# Tonic dopamine regulates cell-type-specific PKA activity and motivated behavior

**DOI:** 10.64898/2026.11.18.767692

**Authors:** Bart Lodder, Tarun Kamath, Isa Hartman, Michelle S. Lopez, Sophie Hoogendoorn, Alloise van Kleef, Kwun Nok Mimi Man, Ximena Opitz-Araya, Michelle Raghubardayal, Joshua Timmins, Roger A.H. Adan, Jonathan Ting, Lin Tian, Bernardo L Sabatini

## Abstract

Dopamine in the striatum fluctuates on fast “phasic” and slow “tonic” timescales. Although disruptions in tonic dopamine have been implicated in the pathophysiology of neuropsychiatric disease, the effect of slow, minutes-long fluctuations in dopamine on neuronal activity and behavior remain unknown. Here we use two-color fluorescence lifetime photometry at high temporal resolution (FLIPR) and calibrated optogenetic and chemogenetic manipulations to determine the impact of tonic dopamine on motivated behavior and downstream signaling *in vivo*. We establish a method for tracking tonic dopamine using FLIPR and determine that dopamine release fluctuates across short (100s of milliseconds) and long (10s of minutes) timescales. We find that dopamine type 1 (D1R) and type 2 (D2R) receptor expressing spiny projection neurons (SPNs) are differentially affected by changes in tonic dopamine in vivo, confirming long-standing hypothesis on tonic dopamine function in the striatum. Finally, we show that accumbal tonic dopamine levels predict and causally mediate an animal’s motivation to work for reward by regulating the rate of reward-motivated actions and engagement in a progressive ratio task. Altogether, we show that tonic dopamine affects downstream signaling and motivated behavior, and establish a new set of methods that allows for careful capture and manipulation of tonic dopamine in vivo.

## Introduction

Striatal dopamine signaling is critical for movement, motivation, and learning, and dysfunction of this system is implicated in neuropsychiatric diseases including addiction, schizophrenia, and Parkinson’s disease ^1,2^. Dopamine neurons display both burst firing and tonic pacemaker firing, which are hypothesized to drive fast (phasic) and slow (tonic) fluctuations in striatal dopamine concentration, respectively ^3,4^. Tonic dopamine levels are also set by molecular factors such as differences in expression of the dopamine transporter and dopamine autoreceptors on dopaminergic axons ^5–7^. Dopamine-deficient mice are hypoactive and unable to independently feed or drink, whereas hyperdopaminergic mice show increased movement, vigor and motivation ^8–10^. Notably, while these animal models have been used to study how tonic dopamine signaling may influence behavior, these models often have alterations in both fast and slow dopamine signaling, limiting the ability to differentiate the behavioral function of each signaling modality.

Fast, phasic dopaminergic fluctuations have been extensively studied and, depending on the striatal subregion, may represent a signal resembling a reward-prediction, action-prediction, or even threat-prediction error ^11–14^. Slow fluctuations in dopamine level are hypothesized to regulate the impact of phasic dopamine on downstream circuits and may correlate with movement, feeding, and motivational state ^15–18^. Computational models suggest that tonic dopamine levels regulate value learning, action selection, and motivation either directly through influencing behavioral processes or indirectly through influencing the phasic dopamine signals and their behavioral effects^19,20^.

Within the striatum, dopamine acts primarily via two G-protein-coupled receptors: the relatively low-affinity dopamine type 1 receptor (D1R) and the relatively high-affinity type 2 receptor (D2R) ^21^. Dopamine binding to D1Rs increases, while dopamine binding to D2Rs decreases, the activity of protein kinase A (PKA), a central kinase that modulates neuronal excitability, plasticity, and gene transcription ^22–26^. In the striatum, these two receptors are expressed on distinct populations of spiny projection neurons (SPNs) ^27^. Previous studies demonstrate that, *in vivo*, phasic dopamine increases activate D1Rs, transiently enhancing PKA activity in D1R-expressing SPNs. In contrast, phasic dopamine increases appear to have little effect on PKA activity in D2R-expressing SPNs which are, instead, derepressed by dopamine unbinding from D2Rs during phasic dopamine dips ^28,29^.

These observations support the hypothesis that *in vivo*, because of differences in ligand affinity, dopamine at normal resting levels minimally occupies D1Rs but nearly maximally binds to D2Rs ^28,30,31^. However, basal dopamine levels have not been measured concurrently with PKA activation, and it is unknown if and how potential slow changes in tonic dopamine dynamically modulate PKA signaling in each SPN class. It has been proposed that changes in tonic dopamine bias phasic dopamine signaling by potentiating absolute phasic increases or decreases, shifting the balance between D1R and D2R sensitivity ^32–34^. An alternative model posits that high-affinity D2Rs detect slow changes in tonic dopamine whereas low-affinity D1Rs detect only phasic dopamine transients, thus mapping a differential temporal filtering of dopamine signals onto the two classes of SPNs ^31,32^. Tonic dopamine may also impact the magnitude of phasic dopamine release via dopamine autoreceptors expressed on dopaminergic axons, thereby primarily affecting the transient changes in PKA activity induced by phasic dopaminergic events ^3,4,35^.

Understanding the function of tonic dopamine in regulating SPNs and controlling behavior is imperative, as tonic dopamine levels are thought to regulate many aspects of behavior, and to be perturbed in many neuropsychiatric diseases ^36,37^. Technical challenges have precluded simultaneous measurement of tonic and phasic dopamine. Methods such as fluorescence intensity-based fiber photometry can measure high-frequency phasic dopamine changes; however, they generally rely on normalization methods and report only relative changes, making measurements of resting dopamine levels and slow changes in dopamine challenging ^38^. Conversely, methods such as microdialysis measure absolute dopamine levels, but are too slow to capture high-frequency phasic dopamine changes, induce significant acute tissue damage, and change the concentration of dopamine during sampling ^39^. Although fast-scan cyclic voltammetry (FSCV), in theory, can provide both absolute measurements and operate over broad temporal scales, challenges in calibration, implementation, and stability have limited its use ^40–42^.

We recently developed fluorescence lifetime photometry at high temporal resolution (FLIPR) and coupled it with novel genetically-encoded dopamine sensors to perform high-frequency, absolute measurements of fast and slow dopamine ^43–47^. Here we employ multi-color FLIPR to measure tonic and phasic dopamine signaling during extended behavioral paradigms, and calibrated optogenetic and chemogenetic manipulations to determine the role and impact of tonic dopamine on D1R-SPN and D2R-SPN PKA signaling and motivated behavior *in vivo*. We find that tonic dopamine fluctuations determine how much an animal works for reward and that these fluctuations meaningfully, but differentially, influence PKA activity in D1R- and D2R-SPNs.

## Results

### Timescales of dopamine signals

To determine the timescales over which dopamine levels fluctuate *in vivo*, we used adeno-associated virus (AAV) to express the genetically encoded dopamine sensor, dLight3.8 or an inactivated dLight3.8 binding mutant (dLight3.8mut3x) in the nucleus accumbens core (NAC) (Fig. 1A). We measured fluorescence lifetime of each in freely moving mice using FLIPR and observed fluctuations in the lifetime of the dLight3.8 sensor at fast (<100ms) and slow (∼minutes) timescales (Fig. 1B,C, S1A). A significant portion of the variance in the fluorescence lifetime occurs at low frequencies corresponding to changes occurring on the order 10s of seconds to minutes (Fig. 1C). Conversely, minimal variance was observed in the lifetime signal of dLight3.8mut3x at these frequencies (Fig. 1C), confirming that the lifetime fluctuations result from interactions at the ligand binding site. Furthermore, ablation of midbrain dopaminergic neurons with 6-OHDA decreased the mean lifetime reported by the dLight3.8 sensor to the same level as observed for dLight3.8mut3x, indicating that basal lifetime levels measured with this approach reflect tonic dopamine levels (Fig. S1B-C).

**Fig. 1:**
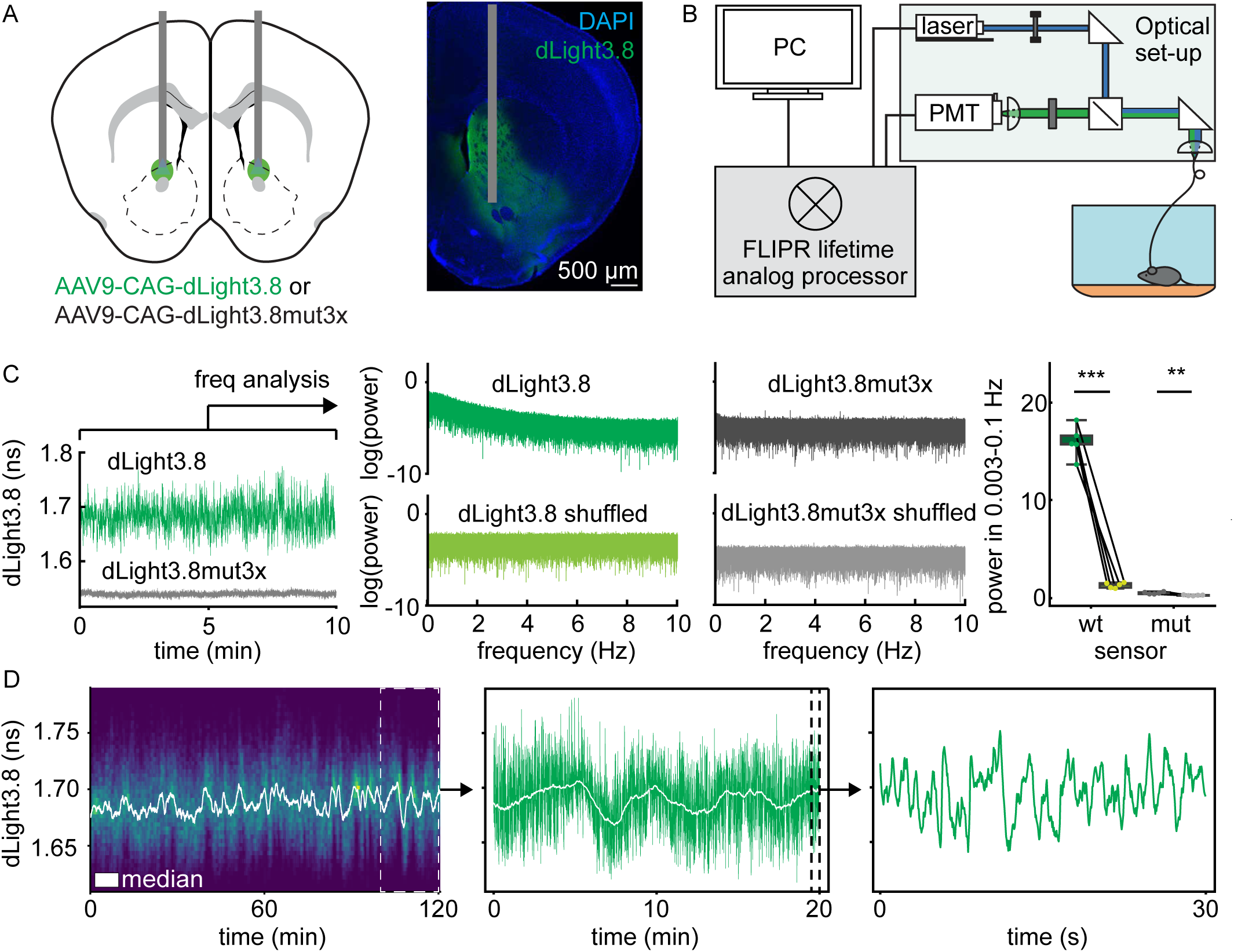
Measuring tonic and phasic dopamine using fluorescence lifetime photometry. **A)** Schematic (*left*) and histology (*right*) showing expression of dLight3.8 or dLight3.8mut and fiber optic (gray) placement in the NAC. **B)** Schematic of FLIPR system connected via fiber optic patch cord to a mouse. **C)** Example timeseries of lifetime measurements of dLight3.8 (green) and dlight3.8mut (gray) (*left*) and the Fourier transforms (*middle*) of the actual or shuffled signals with quantification of power in at low frequencies (*right*; paired sample t test, p=3.4E-5, n=5 dLight3.8 animals; paired sample t-test p=3.3E-3, n=5 dLight3.8x3mut animals). **D)** Example trace of lifetime measurement of dLight3.8 graphed as a pseudocolored rolling histogram with extracted tonic dopamine trace overlaid. inset: an expanded 20 min period. Second inset: an expanded 0.5 min period.

We compared 4 potential methods to extract slow changes in dopamine level over time from dLight3.8 lifetime data: median, mean, iterative Gaussian, and 5th percentile in a rolling 1 minute window (Fig S1D). The moving median, mean, and iterative Gaussian showed high similarity in both lifetime estimate (<5Δps) and correlation (r>0.97) (Fig S1E). The 5^th^ percentile measurement was less correlated and substantially different in lifetime estimate from the median filter-based estimate (>35Δps) (Fig S1D-E). For the following experiments, we used a 1-minute moving median to capture tonic dopamine changes, as it is less sensitive to large phasic variation than a 1-minute moving mean and is a more straightforward analytical method compared to the 1-min moving iterative Gaussian. For the analysis presented below, we operationally define our measure of changes in tonic dopamine levels as the fluctuations in this 1-minute moving median.

### Relationship between tonic dopamine and motivated behavior

To examine the relationship between dynamic changes in tonic dopamine and motivated behavior, we measured changes in dopamine signaling in the NAC of mice performing a progressive ratio task. In this task, mice are trained to perform a specific action (in this case, poking their nose into a specific port) a certain number of times to receive a fixed reward (in this case, a chocolate pellet). With each successful trial and reward retrieval, the number of nosepokes required to receive the subsequent reward increases exponentially. The animal is free to engage or disengage with the task at will. The number of rewards the animal receives before it disengages with the task (in this case, when the animal ceases poking for at least 10 minutes) is known as the “breakpoint”, and is a classically defined measure of the motivational state of an animal (Fig. 2B) ^48,49^.

**Fig. 2:**
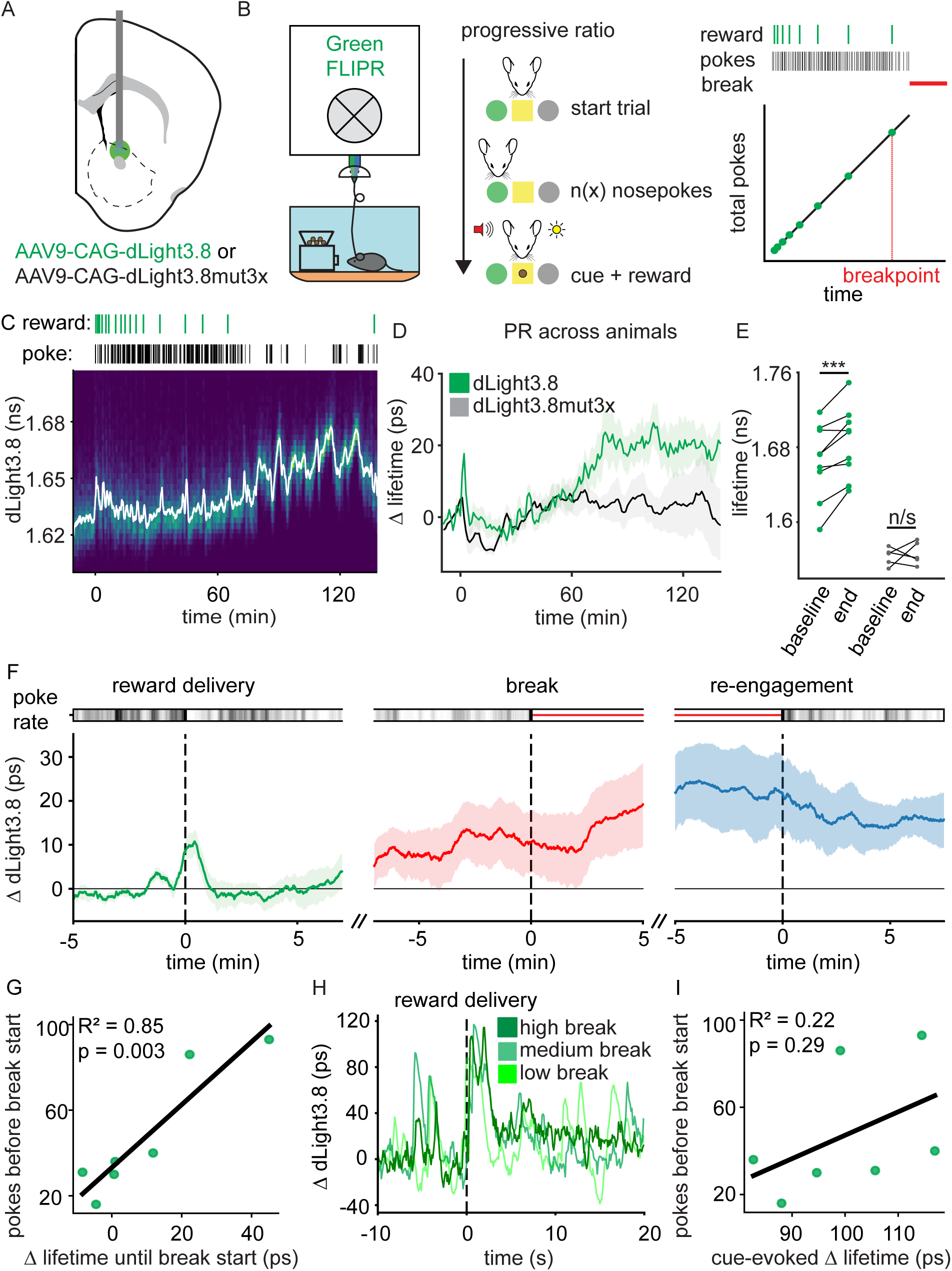
Dopamine fluctuations in the NAC during progressive ratio task. **A)** Schematic of injection/surgery in the NAC. **B)** Schematic of FLIPR setup connected to animal which has access to the FED device (left) and schematic of progressive ratio (PR) task (right). **C)** Example traces of dLight3.8 fluorescence lifetime (*bottom*) during PR task with raster plots (*top*) showing the timing of rewards (green) and reinforced pokes (black). **D)** Average changes in dLight3.8 (green) and dLight3.8x3mut (black) lifetime during PR task. **E)** Quantification of sensor lifetime at the beginning and end of PR task (n=9 dLight3.8 mice, p=0.0027; n=5 dLight3.8x3mut mice, p=0.684, paired sample t-test). **F)** Average change in dLight3.8 lifetime when animals receive the breakpoint pellet (left), when animals cease operantly engaging in the task (middle) and when the animals begin to re-engage after breaking (right) (n=7 dLight3.8 mice). **G)** Correlation between amount an animal works (i.e. number of pokes) in the breakpoint trial until it initially breaks and the change in lifetime during this period. Each point is one animal (n=7 mice, p=0.003). **H)** Example dopamine responses to cue-predicting reward prior to breakpoint in animals at high, medium, or low breakpoint. **I)** Correlation between phasic dopaminergic cue response and pokes leading up to breakpoint (n=7 mice, p=0.292).

We used AAVs to express dLight3.8 or dLight3.8mut in the NAC of mice (Fig. 2A), food-restricted these animals to 90%-95% body weight, and trained them to perform the progressive ratio task. Fluorescence lifetime was recorded for a baseline period prior to the start of the task (∼10 minutes) before the pellet dispensing device was introduced (represented at timepoint t=0), and then continued to be measured for 2.5 hours. Upon introduction of the pellet dispensing device, the fluorescence lifetime increases significantly in dLight3.8 but not dLight3.8mut3x expressing animals (Fig. 2C-D). This change in tonic dopamine level likely reflects a large conditioned response to the pellet-dispensing device. Fluorescence lifetime then consistently increases in dLight3.8 but not dLight3.8mut3x expressing animals during the task, indicating progressive tonic dopamine increases (Fig. 2C-D). Animals showed variability in task performance and dLight3.8 lifetime changes (Fig. 2E, S2). This progressive change in tonic dopamine suggests that as animals must work harder, their tonic dopamine increases. These slow changes in dopamine may reflect the amount of work the animal has invested so far (i.e. sunk cost) or a time-varying internal state which invigorates work (i.e. motivation).

We next specifically investigated how tonic and phasic dopamine change leading up to a breakpoint during the task. First, we isolated those animals which achieved a breakpoint during the task (i.e. spent longer than 10 minutes without operantly working for reward) and specifically looked at the first trial during which the animal broke/disengaged from the task. Immediately prior to the start of these trials, tonic dopamine levels rise transiently upon receipt and consumption of reward before returning to a baseline level (Fig. 2F). Then, during the breakpoint trial, the tonic dopamine level also changes relative to this baseline leading up to disengagement from the task (Fig. 2F). The change in the tonic dopamine level from when the animal receives the previous reward until it disengages is positively correlated with the number of pokes the animal makes during this period, i.e. how much it works for reward (Fig. 2G).

A dopamine phasic increase occurs upon receipt and consumption of the reward in this task, as expected given that the activity of dopaminergic neurons projecting to the NAC is strongly modulated by rewarding stimuli ^50^ (Fig. 2H). We thus isolated the phasic dopamine response to the reward in the trial prior to the breakpoint. In contrast to the predictive power of the tonic dopamine changes, the magnitude of this phasic dopaminergic response is not predictive of the number of pokes the animal subsequently makes before breaking. (Fig. 2I).

In the progressive ratio task, the local temporal reward rate the animal experiences decreases over time as the period between rewards increases. To examine if decreasing reward rate explains changes in tonic dopamine as observed in progressive ratio, we gave mice rewards (chocolate pellets) at varying rates in a non-operant manner (Fig. S3A,B). Phasic fluctuations are observed with each reward (Fig. S3C), but the magnitudes of the tonic or phasic dopamine signals are unaffected by reward rates (Fig. S3C,D). Thus, NAC tonic dopamine increases as animals advance through a progressive ratio task and is correlated with the amount an animal works for a reward before disengaging, indicating that tonic dopamine may reflect or determine the motivation of an animal to work for reward.

### Relationship between dopamine and downstream signaling in behavior

Dopamine fluctuations in the striatum are thought to affect behavior primarily through dopamine Type-1 (D1R) and Type-2 (D2Rs) receptors expressed by SPNs. D1Rs are Gαs-coupled and expressed on D1R-SPNs, whereas D2Rs are Gαi-coupled and expressed on D2R-SPNs. Thus, increases in DA are expected to activate PKA in D1R-SPNs and repress PKA in D2R-SPNs, which subsequently modulates plasticity, excitability and/or gene expression ^21,28,51^. To determine if changes in tonic dopamine are sufficient to affect the biochemical state of each SPN population, we simultaneously measured dopamine levels and PKA activity in D1R- or D2R-SPNs. To do so, we employed FLIM-rdLight1.1, a novel red dopamine lifetime sensor, alongside the green lifetime net PKA activity reporter, FLIM-AKAR ^52–54^.

We first validated that FLIM-rdLight1.1 is utile for measuring dopamine *in vivo*. The absolute lifetime and variance of FLIM-rdLight1.1 are reduced in the presence of a D1R antagonist, confirming that binding at the ligand-binding site is necessary to drive fluorescence lifetime changes (Fig. S4B-C). Optogenetic activation and inhibition of midbrain dopamine neurons increases and decreases sensor lifetime, respectively, confirming that changes in dopamine are sufficient to drive changes in sensor lifetime (Fig. S4E-F). Finally, we benchmarked FLIM-rdLight1.1 against dlight3.8 using a two-color FLIPR system to measure lifetime fluctuations in the two sensors simultaneously (Fig. 3B). We injected animals with AAVs to express dLight3.8 and FLIM-rdLight1.1 in the NAC and implanted a fiber to record fluorescence at the same site (Fig. S4G). dLight3.8 and FLIM-rdLight1.1 lifetime fluctuations are highly correlated and display similar kinetics during spontaneous and behaviorally-induced dopamine fluctuations (Fig. S4H-J).

**Fig. 3:**
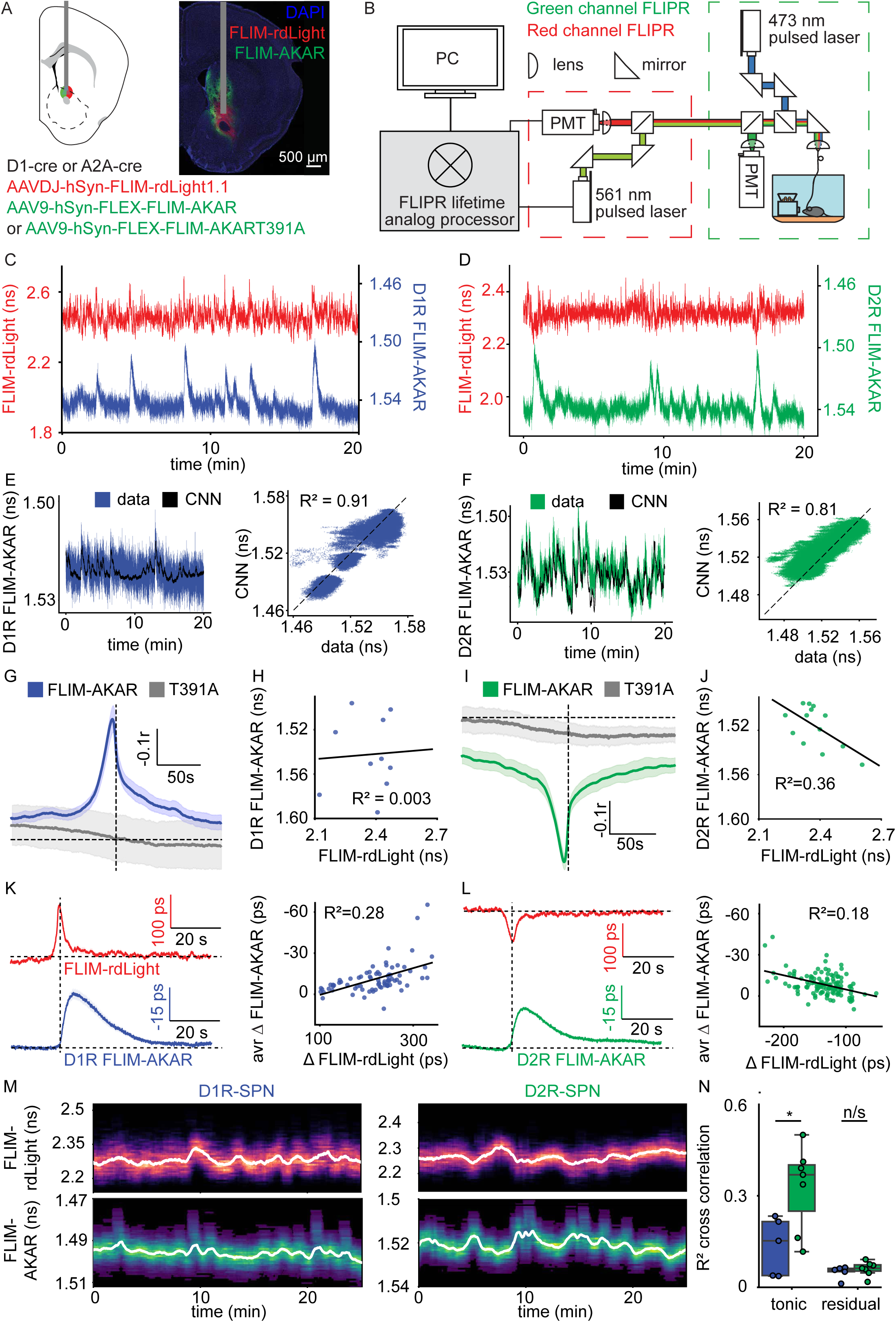
Dopamine levels and PKA activity fluctuations in the NAC. **A)** Schematic (*left)* and histology *(right)* showing expression of FLIM-rdLight1.1 and FLIM-AKAR or FLIM-AKART391A and fiber optic (*gray)* in the NAC. **B)** Schematic of two-color FLIPR setup. **C)** Example recording of FLIM-rdLight1.1 (red) and FLIM-AKAR dynamics in D1R-SPNs (blue). **D)** As in panel B except for D2R-SPN FLIM-AKAR dynamics (green). For all FLIM-AKAR graphs, axis is flipped for ease of interpretation as FLIM-AKAR lifetime is anticorrelated with net PKA activity. **E)** Example trace of D1R-SPN PKA dynamics (blue) and CNN-predicted PKA dynamics from dopamine signal (black, left) and correlation between actual and predicted data (right). **F)** As in panel E, except for D2R-SPN PKA dynamics. **G)** Cross correlation between FLIM-rdLight1.1 and FLIM-AKAR (blue, n=5 mice) and FLIM-rdLight1.1 and FLIM-AKART391A (gray, n=4 mice) in D1R-SPNs. **H)** Correlation between average FLIM-rdLight1.1 level and FLIM-AKAR level in D1R-SPNs across animals (n=10 mice, p=0.877). **I)** As in panel G, but for D2R-SPNs (n=7 FLIM-AKAR mice, n=3 FLIM-AKART391A mice). **J)** As in panel H but for D2R-SPN (n=13 mice, p=0.027). **K)** Average phasic rdLight response (top left) and associated FLIM-AKAR response (bottom left) in D1R-SPNs and regression between magnitude of each response (right, n=77 phasic events, p=5.18x10^-7^). **L)** As in panel K, but for D2R-SPNs (n=122 phasic events, p=6.77x10^-7^). **M)** Example rolling histogram plot of fluctuations in lifetime in FLIM-rdLight1.1 (top) and FLIM-AKAR (bottom) in D1R- (left) and D2R-SPNs (right). Overlying trace represents 1 minute moving median filtered version of each signal. **N)** Comparison of predictive power of different components of FLIM-rdLight1.1 signal for FLIM-AKAR dynamics in D1R- and D2R-SPNs (phasic: p=0.406, tonic: p=0.017; unpaired t-test; n=5 D1R-SPN mice, n=7 D2R-SPN mice).

Therefore, we used AAVs to express FLIM-rdLight1.1 non-specifically and FLIM-AKAR specifically in either D1R- or D2R-SPNS in the NAC (Fig. 3A). FLIM-AKAR fluorescence lifetime is anticorrelated with net PKA activity and is therefore plotted inverted in the figures. We observed that during the progressive ratio task, the relationships between FLIM-rdLight1.1 and FLIM-AKAR lifetimes differed for D1R- and D2R-SPNs, revealing different sensitivity of PKA to slow and fast dopamine changes in each cell type (Fig. 3C-D). To assess whether PKA activity fluctuations can be explained by fluctuations in dopamine, we trained a convolutional neural network to predict FLIM-AKAR lifetime based on FLIM-rdLight1.1 lifetime. This model revealed that NAC dopamine fluctuations are highly predictive of PKA activity in D1R- and D2R-SPNs (Fig. 3E,F).

To interpret which features of dopamine levels and dynamics affect PKA in each cell type, we used linear methods. Grossly, changes in dopamine levels are positively correlated with changes in PKA activity in D1R-SPNs, and anti-correlated with PKA activity in D2R-SPNs - cross correlation revealed that dopamine fluctuations precede PKA activity fluctuations by ∼4.5s (4.2s in D1R-SPNs, and 4.8s in D2R-SPNs) (Fig. 3G, I). Similar correlations are not observed for the lifetimes of FLIM-rdLight1.1 and FLIM-AKART391A, a PKA phosphorylation site mutant version of FLIM-AKAR (Fig. 3G, I). Across mice and recording sites, the absolute average tonic dopamine level is predictive of average PKA activity in D2R-SPNs, but not in D1R-SPNs (Fig. 3H,J), consistent with differential sensitivity of PKA to tonic dopamine in each cell type.

We investigated the contributions of the two modes of dopamine signaling, phasic and tonic, to explaining variance in PKA activity in SPNs. First, to determine the relationship of phasic dopamine with PKA activity, we used an exponential deconvolution algorithm (OASIS) to identify transient SPN PKA events, and analyzed the preceding fluctuation in the dopamine signal^55^. As expected, a phasic increase in dopamine precedes phasic PKA events in D1R-SPNs, whereas a phasic decrease in dopamine precedes phasic PKA events in D2R-SPNs (Fig. 3K-L). Moreover, the magnitude of the preceding phasic dopamine events correlates with the magnitude of the identified PKA event, with opposite signs in D1R- and D2R-SPNs (Fig. 3K-L). Thus, this event-based analysis suggests a causal relationship between phasic dopamine and transient PKA signaling in SPNs.

To analyze the contribution of tonic dopamine signaling to explaining variance in PKA activity in SPNs, we extracted the tonic dopamine signal using the rolling median filter (Fig. 3M). This tonic dopamine signal was cross correlated with the FLIM-AKAR signal in D1R- and D2R-SPNs. Tonic dopamine shifts are more predictive of moment-to-moment fluctuations in net PKA activity in D2R-SPNs than fluctuations in net PKA activity in D1R-SPNs (Fig. 3N). The residual of the signal (i.e. the entire dopamine signal minus the tonic dopamine signal, or only the high frequency component of the dopamine signal) was poorly predictive of PKA activity in D1R-SPNs and D2R-SPNs (Fig. 3N). This analysis thus suggests that D2R-SPNs are more influenced by tonic dopamine changes than D1R-SPNs.

Finally, we determined the relationship between tonic and phasic dopamine and their combined impact on PKA activity. It is possible that at high tonic dopamine levels, further phasic increases in dopamine cannot occur or that at low tonic dopamine levels, further phasic dips cannot occur. This is not the case as tonic dopamine levels are positively correlated with the absolute level of the dopamine phasic events (both increases and decreases) (Fig. S5A-C), indicating that neither increases nor decreases reach a limit and consistent with similar magnitude phasic transients occur on top of varying tonic levels.

Moreover, transient PKA events in D1R-SPNs are more likely to occur in periods of high tonic dopamine, and in D2R-SPNs are more likely to occur in periods of low tonic dopamine (Fig. S5D-F). This suggests that changes in tonic dopamine level influence the magnitudes of phasic dopamine and modulate the probability of triggering a PKA event. Therefore, we examined whether baseline-normalized (i.e. phasic alone) dopamine fluctuations or absolute dopamine fluctuations were more predictive of changes in PKA activity. We found that in both D1R-SPNs and D2R-SPNs, prediction of PKA fluctuations is superior when using the absolute dopamine changes compared to using baseline-normalized changes (Fig. S5G-N). Together, this suggests that tonic dopamine influences the absolute magnitude of phasic dopamine changes, which in turn shapes the probability and magnitude of PKA events in D1R- and D2R-SPNs.

### Tonic dopamine differentially impacts D1R- and D2R-SPN PKA signaling

Although the results above suggest a relationship between tonic and phasic dopamine and D1R- and D2R-SPN PKA signaling, external variables, such as other neuromodulators coinciding with dopaminergic fluctuations, may confound the analyses. Therefore, we directly manipulated tonic and phasic dopamine and measured effects on PKA activity in SPNs. We used AAVs to express somatically-targeted BiPoles (a bidirectional opsin consisting of a linked excitatory opsin, Chrimson, and an inhibitory opsin, gtACR2) in ventral tegmental area (VTA) dopamine neurons (DANs) and implanted a fiber above the midbrain injection site ^56^. Additionally, we used AAVs to express FLIM-rdLight1.1 and FLIM-AKAR in the NAC, the latter specifically in D1R- or D2R-SPNs, and placed an optical fiber above this injection site to monitor striatal dopamine levels and cell-type specific PKA activity (Fig. 4A). We calibrated phasic and tonic manipulations of VTA DANs to evoke a range of dopaminergic responses in freely moving mice (Fig. 4B,C) similar to those previously observed (Fig 3). We measured the effects of these manipulations on dopamine levels and PKA activity in D1R- and D2R-SPNs.

**Fig. 4:**
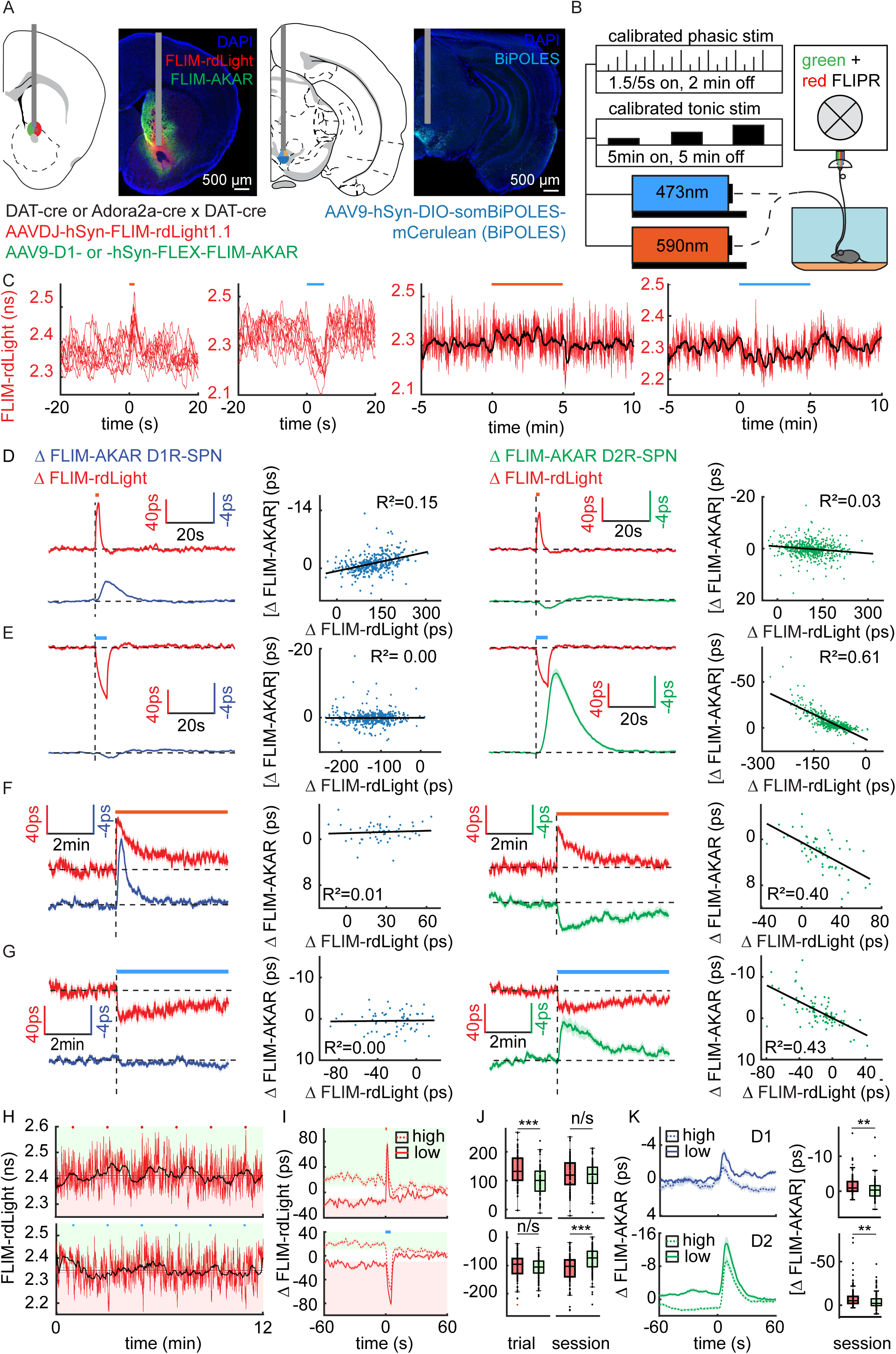
Optogenetically induced phasic and tonic dopamine fluctuations and effects on PKA activity in striatal SPNs. **A)** Schematic (*left)* and histology (*right*) showing expression of FLIM-rdLight1.1 and FLIM-AKAR in the NAC and somBiPoLes in the VTA with fiber optics (*gray*) for recording and excitation of opsin, respectively. **B)** Schematic of two-color FLIPR setup with integrated optogenetic stimulation. **C)** Example optogenetically-induced phasic increases and decreases in dopamine (left) and optogenetically-induced tonic increases and decreases in dopamine (right). Overlying black line represents a 20s moving median filter. **D)** Left: average optogenetically-induced phasic increase in rdLight signal and associated D1R-SPN FLIM-AKAR dynamic, with regression between magnitude of evoked rdLight change and associated FLIM-AKAR change in D1R-SPN (n=367 stimulations, across n=15 sessions, in n=6 sites across N=5 animals; p=7.61x10^-15^). Right: as on left side except for D2R-SPNs (n=546 stimulations across n=21 sessions, in n=6 sites across N=4 animals; p=1.78x10^-4^). **E)** As in panel D except for changes seen with optogenetically-induced phasic decreases in rdLight signal (D1R-SPN: n=450 stimulations, across n=16 sessions, in n=6 sites across N=5 animals, p=0.94; D2R-SPN: n=511 stimulations across n=18 sessions, in n=6 sites across N=4 animals, p=2.09x10^-107^). **F)** Left: average optogenetically induced tonic increase in rdLight signal and associated D1R-SPN FLIM-AKAR dynamics across the five-minute period (analysis only performed on last two minutes). Regression between median evoked rdLight change and average associated change in FLIM-AKAR signal in D1R-SPNs (n=53 stimulations across n=12 sessions, in n=6 sites across N=5 animals, p=0.60). Right: as on left side except for D2R-SPNs (n=59 stimulations across n=10 sessions, in n=6 sites, across N=4 animals, p=6.62x10^-8^). **G)** As in panel F except for changes seen with optogenetically-induced tonic decreases in rdLight signal (D1R-SPN: n=59 stimulations across n=10 sessions, in n=5 sites across N=5 animals, p=0.855; D2R-SPN: n=85 stimulations across n=15 sessions, in n=6 sites across N=4 animals, p=8.22x10^-12^). **H)** Example trace of optogenetically-evoked phasic increases (top) and decreases (bottom) in rdLight lifetime at different periods of low (red) versus high (green) tonic dopamine. **I)** Average optogenetically-evoked phasic increase (top) and decrease (bottom) in dopamine signal at periods of high (green) or low (red) tonic dopamine. Dopamine signal is normalized to the entire session average. **J)** Comparison of magnitude of phasic dopamine changes normalized either on a per stimulation (i.e. trial, left; top: n=84 stimulations, across n=15 sessions, in n=6 sites across N=5 mice, p=4.127E-06, bottom: n=109 stimulations across n=15 sessions, in n=6 sites across N=5 mice, p=0.157; unpaired t-test) or per session (right; top: 0.635, bottom: p=1.27x10^-6^; unpaired t-test) basis. **K)** Associated change in D1R-SPN (top) and D2R-SPN (bottom) PKA change with optogenetic phasic increases and decreases in dopamine, respectively at higher (green) or lower (red) tonic dopamine levels. Right: quantification of these PKA changes (top: n=126 stimulations, across n = 18 sessions, in n=6 sites across N=4 mice, p=0.0084, bottom: n=136 stimulations, across n=18 sessions, in n=6 sites, across N=4 mice, p=0.0058; unpaired t-test).

In line with prior studies and our own results, phasic increases in dopamine increase PKA activity in D1R-SPNs, but minimally affect D2R-SPN PKA activity (Fig. 4D) ^57^. Conversely, phasic decreases in dopamine increase D2R-SPN PKA activity, but minimally affect D1R-SPN PKA activity (Fig. 4E). PKA activity responses in each cell type scale with the magnitude of the phasic dopaminergic response (Fig. 4D,E).

To investigate the impact of tonic dopamine changes on striatal cell-type specific PKA activity, we optogenetically manipulated dopamine neurons for five minutes (Fig. 4C) and analyzed dopamine and PKA activity changes in the final two minutes to avoid confounding by potential effects of phasic dopamine changes at the beginning of the optogenetic stimulus. Optogenetically-evoked changes in tonic dopamine bidirectionally affect PKA activity in D2R-SPNs, but not in D1R-SPNs (Fig. 4F,G, S6). The impact on D2R-SPN PKA scales linearly with the magnitude of increases and decrease in tonic dopamine, as opposed to the phasic changes which only scale with optogenetically-evoked decreases in dopamine, but not increases.

The optogenetically-evoked phasic dopamine transients occurred by chance during periods of higher or lower tonic dopamine levels. We examined how this naturally occurring variance in tonic dopamine influences phasic dopamine transients and SPN PKA activity. For each dopamine recording, we normalized the data to the average tonic dopamine level across the session, allowing for comparisons within a given session of tonic and phasic dopamine changes. We categorized optogenetically-evoked phasic dopaminergic events based on whether they occurred during periods of high (>10ps above session baseline) or low (>10ps below session baseline) tonic dopamine (Fig. 4H).

Optogenetic stimulation of midbrain DANs evokes significantly larger phasic dopamine increases at lower tonic dopamine levels compared to the same stimulation at higher tonic levels (Fig 4I,J). The D1R-SPN PKA response triggered by the smaller phasic dopamine event on a higher tonic dopamine level is smaller than the PKA response which is triggered by a larger phasic dopamine event on a lower tonic dopamine level (Fig. 4K). Therefore, while slow dopamine shifts do not directly affect PKA activity in D1R-SPNs, the tonic dopamine level indirectly influences transient PKA activity in D1R-SPNs by modulating the magnitude of phasic dopamine increases.

Optogenetic inhibition of midbrain DANs evokes a similar magnitude phasic dopamine decrease at high and low tonic dopamine levels (Fig. 4I,J). Thus, the optogenetically-evoked dopamine minimum varied as a function of the tonic dopamine level. The D2R-SPN PKA response triggered by a phasic dopamine dip on a higher tonic dopamine level is smaller than that which is triggered by a phasic dopamine dip on a lower tonic dopamine level (Fig. 4K). Thus D2R-SPN PKA events triggered by phasic dopamine dips scale with the absolute dopamine level rather than purely the phasic magnitude. Overall, we thus find that slow changes in dopamine differentially effect PKA signaling in each striatal cell type - in D2R-SPNs, slow dopamine changes directly influence moment-to-moment PKA activity, while in D1R-SPNs, tonic dopamine levels indirectly influence PKA activity by modulating the magnitude of phasic dopamine increases.

### Tonic dopamine levels determine motivation to work for reward

Thus, tonic dopamine is correlated with behavior and motivational variables, and these tonic dopamine changes have a meaningful influence on downstream neural circuitry. However, it is still unclear from the aforementioned results whether slow changes in dopamine are reflective of changes in motivational state or actually determine an animal’s motivation. Thus, to determine if these tonic dopamine changes impact behavior, we manipulated tonic dopamine during the progressive ratio task and measured the effects on motivation as reported by the breakpoint. To manipulate tonic dopamine for extended periods, we employed a chemogenetic approach and used AAVs to express excitatory (Gq-coupled) or inhibitory (Gi-coupled) designer receptors exclusively activated by designer drugs (DREADDs) in VTA DANs ^58^. The effects of these manipulations on dopamine levels were monitored via dLight3.8 lifetime photometry in the NAC (Fig. 5A).

**Fig. 5:**
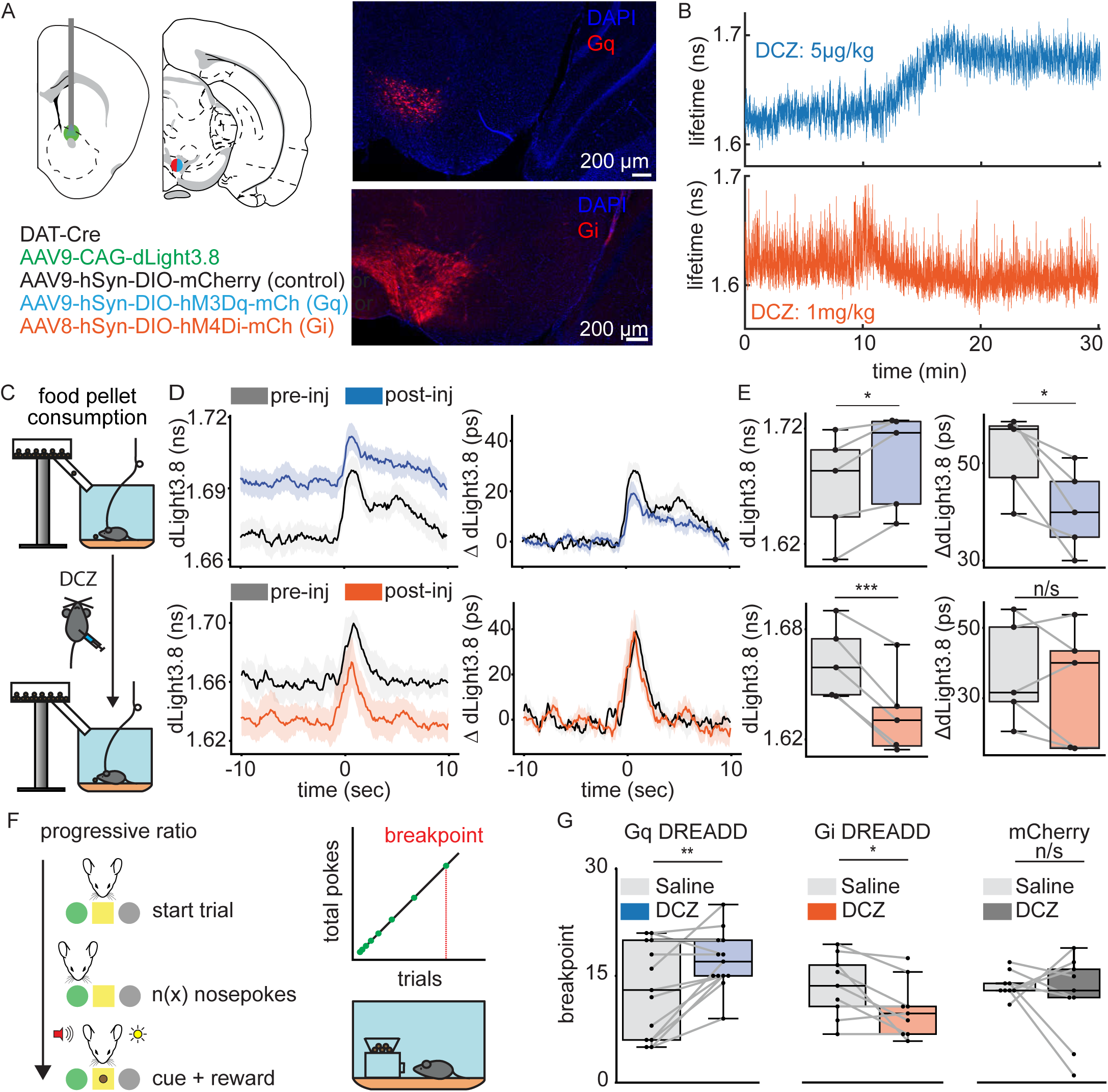
Effect of tonic dopamine changes on motivation to work for reward. **A)** Schematic of injection/surgery in the NAC and VTA and associated histological section of Gq (top) and Gi (bottom) DREADD spread. **B)** Example dLight3.8 lifetime changes associated with DCZ injection in animals expressing either Gq (top) or Gi (bottom) DREADDs in VTA DANs. **C)** Schematic of experiment to test if evoked phasic dopamine events are affected by DREADD mediated manipulation of tonic dopamine levels. **D)** Phasic dopamine response evoked by food pellet consumption before (gray) and after (blue/orange) injection of DCZ in animals expressing Gq (top) or Gi (bottom) DREADDs either shown as absolute (left) or normalized (right) traces. **E)** Tonic (left) and phasic (right) dopamine changes during the task in mice expressing Gq (top, n=5 mice; tonic: p=0.0145, phasic: p=0.0147; paired t-test) or Gi (bottom, n=5 mice; tonic: p=0.00065, phasic: p=0.447; paired t-test) DREADDs. **F)** Schematic of progressive ratio task with measurement of breakpoint in mice. **G)** Comparison of breakpoint in animals injected with either saline (grey) or DCZ (blue/orange/dark grey) in mice expressing Gq DREADDs (n=13 mice, p=0.0078, paired t-test), Gi DREADDs (n=9 mice, p=0.029, paired t-test) or mCherry (n=9 mice, p=0.459, paired t-test) in VTA DANs. All comparisons are done on a per-animal basis (comparing the same animal with an injection of saline vs DCZ).

We used IP administration of deschloroclozapine (DCZ) to activate the DREADDs ^59^. We tested multiple doses of DCZ and found that at higher doses, DCZ-induced activation of Gq DREADDs in DANs substantially diminished high-frequency dopaminergic activity and led to supraphysiological, disordered, and inconsistent patterns of dopamine release (Fig. S7A). We thus calibrated the DCZ dosage to elicit a tonic dopamine change that minimally affects spontaneous phasic activity (Fig. 5B) and does not change tonic dopamine levels in the absence of DREADDs (Fig. S7B). To investigate whether evoked phasic dopamine release was affected by the tonic dopamine manipulation, we food-restricted mice and gave them chocolate pellets, a rewarding stimulus known to evoke a phasic dopamine response, before and after manipulation of the tonic dopamine level and measured the magnitude of the phasic dopamine response (Fig 5C). Decreasing tonic dopamine using DREADDs does not change the phasic dopamine response to a food pellet reward (Fig 5D-E), whereas increasing tonic dopamine using DREADDs results in a small, but significant, decrease in the phasic dopamine response to the food pellet (Fig 5D-E).

Mice were trained and tested in the progressive ratio task and received saline (control) or DCZ on alternating days (Fig. 5F). Increasing tonic dopamine increases the breakpoint and poke rate, whereas decreasing tonic dopamine decreases the breakpoint and poke rate (Fig 5G, S7D). These effects in progressive ratio tasks are interpreted as, respectively, increasing and decreasing the animal’s motivation to work for reward. DCZ alone in the absence of DREADDs does not significantly change an animal’s breakpoint nor its poke rate, i.e. does not change its motivation to work for reward (Fig 5G, S7D). Thus, tonic dopamine levels drive an animal’s motivation to work for reward.

## Discussion

In this study, we apply fluorescence lifetime photometry at high temporal resolution (FLIPR) with novel genetically-encoded lifetime sensors and calibrated optogenetic and chemogenetic manipulations to investigate how tonic dopamine signaling affects striatal circuitry and animal behavior. We find that dopamine fluctuates on relatively short (sub-second) and relatively long (seconds to minutes) timescales. Importantly, we find that tonic dopamine is not static, but rather is dynamically regulated. Tonic dopamine levels in turn modulate an animal’s motivation to work for reward. Finally, these slow changes in dopamine selectively engage and influence PKA activity in D2R-SPNs and regulate the influence that phasic dopamine changes have on PKA activity in D1R-SPNs. We thus delineate how slow dopaminergic changes regulate striatal plasticity *in vivo* and an animal’s motivation to work for reward.

### Tonic dopamine modulation of motivation to work for reward

The behavioral influence of slow versus fast changes in striatal dopamine levels has been difficult to study, as technologies used to record dopamine *in vivo* are typically optimal for measuring changes either on relatively short or long timescales, but not both ^38–40,42^. We employed FLIPR to measure both fast and slow modulation of dopamine levels in the same animal. We also exploit the stability and robustness of FLIPR, which does not suffer from many artifacts which plague intensity-based methods (ex. photobleaching, differences in sensor expression) ^60^ to validate experimental manipulations of tonic dopamine signaling.

Tonic dopamine has been hypothesized to regulate motivational behavior, based on both experimental and computational models ^16,17,61^. In the progressive ratio task, we find that changes in tonic dopamine are correlated with how much animals work before disengaging, in line with these prior models ^16,62,63^. To determine causality, we chemogenetically manipulated tonic dopamine levels, and found that tonic dopamine levels regulate an animal’s motivation to work for reward. This result is concordant with prior studies that employed chemogenetic and pharmacological manipulations of the dopaminergic system (but did so without measuring the effects on tonic and phasic dopamine) and reported changes in an animal’s motivation to work for reward ^64,65^.

We tailored our manipulation method to selectively affect tonic dopamine with minimal effects on phasic signaling by measuring their effects with FLIPR. This was necessary as higher doses of DCZ used to activate excitatory DREADDs expressed in dopaminergic neurons can affect tonic dopamine levels while drastically suppressing phasic dopamine (thus limiting interpretability of the manipulation) and cause supraphysiological changes in tonic dopamine (which may cause different behavioral effects from physiological release patterns) ^66–68^. Future studies should thus ensure pairing such manipulations with recordings of tonic and phasic dopamine to ensure specificity of manipulation.

Even with this calibrated approach, however, chemogenetically-driven increases in tonic dopamine do slightly decrease phasic dopamine increases (Fig. 5C). This is unlikely an artifact of the manipulation method, as a similar anti-correlation between tonic and phasic dopamine is present when optogenetically stimulating DANs at different tonic dopamine levels (Fig. 4I,J). This suggests that high tonic dopamine levels inhibit phasic dopamine increases (as has long been hypothesized), potentially through changes in vesicular availability or through activation of D2 auto-receptors that inhibit release from dopaminergic axons ^3,6,32,35^. This effect of our manipulation may have influences on the motivational phenotype observed as prior studies have found that decreased phasic dopamine increases can increase an animal’s motivation to work for reward ^69^. Future work could thus manipulate phasic dopamine signals in the progressive ratio task and other behavioral paradigms for studying motivation while measuring tonic dopamine levels.

### Tonic dopamine modulation of biochemical signaling pathways in striatal SPNs

Dopamine putatively influences neural circuitry and animal behavior by modulating the PKA activity and therefore plasticity of striatal SPNs through binding to D1R and D2Rs ^21–26^. Phasic dopamine signals differentially affect SPNs, whereby phasic increases and decreases in dopamine lead to changes in PKA activity in D1R- and D2R-SPNs, respectively. While several models have been put forward on how tonic and phasic dopamine could interact to affect D1R- and D2R-SPNs PKA activity, these models have been difficult to test *in vivo* due to technological limitations in measuring tonic and phasic dopamine ^3,4,32,33^. There may be additional rapid effects of tonic and phasic dopamine fluctuations on the acute activity and subsequent synaptic transmission of SPNs that we do not study here ^70^.

We find that fast dopamine changes cause transient changes in PKA activity in both D1R- and D2R-SPNs. Conversely, slow dopamine shifts alter moment-to-moment PKA activity in D2R-SPNs but not in D1R-SPNs, potentially because the relatively high-affinity D2Rs are more sensitive to these slow changes compared to the relatively low-affinity D1Rs ^30,31^. Thus, these two classes of SPNs separate dopamine signaling in both the sign of the phasic dopaminergic signal and in the frequency component of the dopamine signal, as suggested by previous longstanding models ^31,32^. It is of course possible that slow dopamine shifts polysynaptically influence SPN PKA activity through action on dopamine receptors on non-SPN striatal cells (ex. by binding to D2Rs on cholinergic interneurons and thereby change acetylcholine release) ^71,72^.

In terms of how tonic and phasic dopamine interact to influence SPN PKA activity, we find that tonic dopamine levels alter the magnitude of phasic dopamine increases, but not phasic dopamine decreases. The magnitude of a D1R-SPN PKA event triggered by a phasic increase in dopamine is a function of the magnitude of the phasic event. Thus, paradoxically, higher tonic dopamine levels lead to smaller D1R-SPN PKA events, as the higher tonic dopamine level suppresses phasic release. This is discrepant from our finding that higher tonic dopamine levels are correlated with higher absolute dopamine events (Fig. S5A-C) and a greater probability of a D1R-SPN PKA event (Fig. S5D-F). This discrepancy may be due to fluctuations in other striatal neuromodulators which occur during periods of high tonic dopamine and thus potentiate phasic dopamine events and D1R-SPN PKA activity ^73^. Future work could pair recordings of tonic dopamine shifts with recordings of other striatal neuromodulators to investigate how these neuromodulators co-vary and jointly influence D1R-SPNs PKA activity *in vivo*.

Since tonic dopamine level does not alter the magnitude of phasic dopamine dips, the same phasic dopamine dip on a lower baseline leads to a lower absolute dopamine level, and in turn a larger D2R-SPN PKA event. Thus, changes in tonic dopamine level do alter the magnitude of phasic dopamine induced transient D2R-SPN PKA events. This result is in line with prior models which posit that tonic dopamine level biases the effects of phasic dopaminergic changes on downstream striatal circuitry ^22,33^.

Of course, these measurements aggregate tonic and phasic dopamine changes across many neurons. Future studies should examine the role of tonic dopamine in regulating phasic dopamine and downstream signaling at the single-neuron level using methods such as 2-photon fluorescence lifetime microscopy to discern whether these bulk relationships hold on a per-cell basis. Additionally, future studies should investigate the influence these slow changes in dopamine and resulting changes in PKA activity have on the activity of striatal SPNs *in vivo*.

### Different frequencies of dopamine signaling

The definitions of tonic and phasic dopamine and their associated timescales were initially based on dopamine neuron firing rates and later the sampling rate of dopamine measurement methods ^3,4^. Although here we have primarily studied two timescales of dopamine signaling: slow, “tonic” dopamine changes occurring on 1-minute scales, and fast, “phasic” dopamine changes occurring on milliseconds to seconds timescales, there is significant variance in the dopamine signal across a large frequency spectrum. Thus, the binary definition of tonic and phasic dopamine will need to be refined to encompass these different timescales. Further work could also delineate circuit-level and behavioral consequences of these many different frequencies of dopamine signaling.

Altogether, this work uses pioneering measurement and manipulation methodologies to discern the role of tonic dopamine in motivated behavior and striatal neuron plasticity-controlling PKA dynamics *in vivo*. These same approaches can be used to study tonic dopamine in other striatal subregions, and in other behavioral paradigms, such as during spontaneous movement and in associative learning. Finally, given that tonic dopamine is hypothesized to be central to the pathophysiology of several neuropsychiatric diseases and given that several medications are thought to have therapeutic efficacy by changing tonic dopamine levels, the increased understanding of tonic dopamine and downstream signaling presented in this and subsequent work will pave the way for improved understanding of neuropsychiatric diseases and potential subsequent drug development.

## Author Contributions and Acknowledgements

All experiments were designed by B.L, T.K., R.A.H.A., and B.L.S. Experiments were conducted by B.L., T.K., I.H., M.S.L., S.H., A.v.K., M.R., and J.T. Data was analyzed by B.L. and T.K. The FLIM-rdLight1.1 sensor was designed and constructed by K.N.M.M. and L.T. The enhancer virus for D1R-SPNs was designed and constructed by X.O. and J.T. All authors reviewed the manuscript. We thank all Sabatini lab members for discussions, Bastijn van den Boom, Yang Ge, Shun Li, Sanika Ganesh, and Allison Girasole for evaluating the manuscript, Ofer Mazor of the Harvard Medical School Research Instrumentation Core for building the box housing for the FLIPR system, Adrian Coscia for assisting in the design of dLight3.8x3mut and the Harvard Medical School Machine shop for designing and building parts of the behavior system. This work is supported by grants to B.L.S. (NIH R35NS137336, R01MH140249), B.L. (PhD fellowship, Boehringer Ingelheim Fonds), T.K. (NIH F30DA061675, T32AG51108, T32GM144273), J.T.T. (NIH UF1MH128339), and L.T. (NIH U19NS123719, UM1MH136462).

## Material and methods

### Mice

Experiments were conducted in accordance with protocols approved by the Harvard Standing Committee on Animal Care following principles described in the US National Institutes of Health Guide for the Care and Use of Laboratory Animals. The following mouse lines were used: C57BL/6J (The Jackson Laboratory, 000664), *Adora2a-Cre* (B6.FVB(Cg)-Tg(Adora2a-cre)KG139Gsat/Mmucd, 036158-UCD), *Drd1a-cre* (B6.FVB(Cg)-Tg(Drd1-cre)EY262Gsat/Mmucd, 030989-UCD) (both were originally acquired from MMRRC UC Davis), *DAT-ires-*Cre (The Jackson Laboratory, 006660). Mice of the *Adora2a-crexDAT-ires-Cre* genotype were bred in-house. Female and male mice were used in approximately equal proportions. Ages of mice used in experiments were typically between 2-4 months. During behavioral experimentation, mice were group housed unless with a head-plate or during periods of food-restriction. All mice were housed on a reversed 12hr dark/12hr light cycle, and experimental procedures were performed between 9:00 and 19:00 each day, and animal behavior was assessed at roughly the same time each day.

### Viruses

Serotypes 8, 9, PHP.eB, DJ, and DJ/9 recombinant adeno-associated viruses (AAVs) were injected intracranially to express constructs of interest in a Cre recombinase-dependent or recombinase-independent manner. In all cases, 300nl of virus was injected at a concentration of 10^11^ to 10^13^ genomic copies per ml. AAVs were packaged in-house or by commercial vector core facilities (Janelia Vector Core and UNC Vector Core) and stored at −80°C. The following AAVs were used:

AAV-CAG-dLight3.8 (UNC Neurotools)

AAV-CAG-dLight3.8x3mut (available upon request to Dr. Bernardo Sabatini)

AAVDJ-hSyn-FLIM-rdLight1.1 (available upon request to Dr. Lin Tian)

AAV-hSyn-FLEX-FLIM-AKAR (Addgene #60445)

AAV-hSyn-FLEX-FLIM-AKART391A (Addgene #60446)

AAV-hSyn-FLEX-somBiPOLES-mCerulean (Addgene #154945)

AAV-AiE0779m_3xC2-minBG-FLIM-AKAR-WPRE3-BGHpA (available upon request to Dr. Jonathan Ting)

AAV-hSyn-DIO-hM3D(G)q-mCherry (Addgene #44361)

AAV-hSyn-DIO-mCherry (Addgene #50459)

AAV-hSyn-DIO-hM4D(G)i-mCherry (Addgene #44362)

### Stereotaxic surgical procedures

For all surgical procedures, one day before surgery, mice were given oral carprofen *ad libitum*. Mice were initially and persistently anesthetized with inhaled isoflurane (1-3%) throughout the procedure. Under the stereotaxic frame (David Kopf Instruments), the animal’s skull was exposed and leveled. A small burr hole was made using a drill based on bregma and lambda skull suture lines at the sites for viral injection. After burr hole creation, virus or chemical was injected at a rate of either 50 nL/min (NAC, VTA) or 100 nL/min (MFB) using a syringe pump (Harvard Apparatus, #883015). The pipette was allowed to rest at the injection site and 100 micrometers above the injection site for at least 5 minutes before slow withdrawal. The following titers were injected for each virus: AAV9-CAG-dLight3.8 (7.4-10E12 gc/mL), AAV9-hSyn-dLight3.8mut3x (1-10E12 gc/ml), AAV9-hSyn-FLEX-FLIM-AKAR (0.9-5.6E12 gc/mL), AAV-hSyn-FLEX-FLIM-AKART391A (1.5E12 gc/mL), AAV-AiE0779m_3xC2-minBG-FLIM-AKAR-WPRE3-BGHpA (8E12 gc/mL), AAVDJ-hSyn-FLIM-rdLight1.1 (7.2E12 gc/mL), AAV9-hSyn-FLEX-somBiPOLES-mCerulean (5E12 gc/mL), AAV9-hSyn-DIO-hM3Dq-mCherry (0.7-2.1E12 gc/mL), AAV8-hSyn-DIO-mCherry (2.1E12) and AAV8-hSyn-DIO-hM4Di-mCherry (2.1E13).

The following coordinates were used to inject in the region of interest (anteroposterior (AP), medio-lateral (ML) relative to bregma; dorsoventral (DV) from brain surface, coordinates are in mm):

VTA: -3.1 AP, +/−0.5 ML, -4.1 DV

NAC: +1.4 AP, +/−1.3 ML, -3.85 DV

MFB: -1.0 AP, +/- 1.0 ML, -4.9 DV

For AAV or chemical injections alone, the skin was then sutured shut. For experiments requiring a fiber optic implant, the skull was lightly scored with a razor blade and the fiber optic implant was placed 100 μm above the injection site. For optogenetic experiments, a dual optic fiber was placed (DFC_200/245-0.37_5mm_DF1.0_FLT Dual Fiberoptic Cannula, Doric Lenses). For FLIPR experiments, an mono optical fiber implant (MFC_200/230-0.37_4.5mm_MF1.25_FLT or MFC_200/230-0.37_3mm_MF1.25 mono fiber optic cannula, Doric Lenses) was placed. For photometry experiments alone, recordings began at least three weeks postoperatively. For DREADD and optogenetic manipulation experiments, manipulations and recordings began at least four weeks postoperatively.

### Chemical ablation of dopaminergic neurons

To ablate dopaminergic neurons we employed a protocol as in prior studies ^13^. First we prepared a solution of 14.25 mg desipramine (Sigma-Aldrich, D3900-1G)/3.1 mg pargyline (Sigma-Aldrich, P8013-500MG) dissolved in 5 mL of water. This cocktail increases the selectivity of the chemical ablation of 6-OHDA for dopaminergic neurons and reduces effects on noradrenergic neurons (though these minimally project to the NAC). The desipramine/pargyline solution was injected IP to animals at 10 mg/kg about 30 minutes prior to ablation procedure. 6-OHDA was used to ablate dopamine neurons and was prepared at 10 mg/mL of 6-OHDA (Sigma-Aldrich, H116-5MG) in 0.2% ascorbic acid in saline (Sigma-Aldrich, PHR1008-25MG). Due to photosensitivity, the 6-OHDA mixture was prepared and injected in the dark. 1 µL was injected unilaterally at the MFB (following the abovementioned stereotaxic surgical procedures).

### Behavior

For all experiments, mice were habituated to handling, patch cord attachment, and/or head fixation prior to experimentation. For experiments involving lifetime photometry recordings, mice were connected to the FLIPR system by attaching the optical patch cord to the fiber implant with a ceramic ferrule sleeve (ADAL1-5, Thorlabs). Mice were then allowed to rest in the experimental chamber after attachment to the patch cord for five minutes before beginning recordings.

#### Spontaneous tonic and phasic dopamine behavioral task

To investigate spontaneous changes in fluorescence lifetime putatively representing shifts in tonic and phasic dopamine (in Figure 1 and S1), mice were placed in an 8-by-8-inch experimental chamber. Mouse movement was captured using a camera (FL3-U3-13E4M, PointGrey) operated through Bonsai-Rx software (v2.5.1) ^74^. Animals were allowed to explore the chamber for at least two hours before being removed. Animals were not food-restricted prior to this experiment.

#### Progressive Ratio Behavioral Task

For the progressive ratio task, mice were initially food-restricted overnight and were given 10-14 chocolate-flavored pellets (F05301, Bio Serv). Throughout training and experimental testing, mice were maintained at 90-95% of their original body weight. Mice were trained to perform progressive ratio using a feeding device (FED3, LABmaker) ^75^. Mice were trained on consecutive days for three hours per session at roughly the same time each day. On day 1, chocolate pellets were dispensed from the device freely, with each pellet delivery triggered upon retrieval of the prior pellet. On days 2 and 3, mice were trained to operantly work for pellet rewards, in which they had to poke their nose either one (FR1) or three (FR3) times, respectively, into the left active noseport (the right noseport served as a control) to receive a pellet. On days 4-6, mice were trained on the progressive ratio task wherein mice had to progressively increase the number of nose pokes on the left port to trigger pellet delivery, according to the following formula:

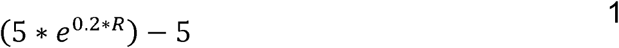

Where R represents the number of rewards the mouse has already received. Training was repeated if mice did not consume a significant number of chocolate-flavored pellets (F05301, Bio Serv) during free feeding, FR1 or FR3 (pellets <40). After day 6, mice were measured using single- or dual-color FLIPR while performing progressive ratio. Behavior was logged using the FED3.0 device and aligned to FLIPR data using video data captured with Bonsai-Rx (v2.5.1).

#### Reward Rate Behavioral Task

To study dopamine signals evoked by pellet consumption, mice were food-restricted overnight and habituated to 10-14 chocolate pellets. Mice were placed in an 8-by-8-inch experimental chamber and were measured using FLIPR. Food pellets were delivered via a pellet-delivery system positioned above the chamber. Feeding behavior was captured using a camera (FL3-U3-13E4M, PointGrey) controlled through Bonsai software (v2.5.1).

#### Pharmacological manipulations

For D1R antagonist SCH23390 hydrochloride (10mg/kg, #0925, Tocris) or saline (00409-4888-10, Hospira) injection, mice were placed in a 6.5 by 14.5-inch experimental chamber and injected intraperitoneally (IP) using insulin injection needles (#1612712B, Sol-M). IP injections were timed by the experimenter and captured using a camera for data alignment.

### Fluorescence lifetime photometry

Fluorescence lifetime photometry was performed as described previously. In short, the two-color FLIPR system consists of a combined green-and-red fluorescence lifetime photometry system connected to FLIPR analog computing units for high-speed lifetime computation. A blue 473nm picosecond pulsed (BDS-SM-473-FBC, Becker & Hickl) was power modulated using an AOM (MTS110-A3-VIS, AA optoelectronic) and a rotary neutral density filter (NDC-50C-4M-A, Thorlabs), reflected on a 488LP dichroic beamsplitter (Di02-R488-25x36, Semrock, mounted in DFM1-P01, Thorlabs) and focused into a patchcord (MFP_200/220/900-0.37_2m_FCM-MF1.25(F)_LAF, Doric). Green fluorescence exiting from the patchcord passes through the 488LP dichroic beamsplitter and was subsequently reflected off a 552LP dichroic beamsplitter (FF552-Di02-25x36, IDEX), filtered using an interference filter (FF01-525/35-25, Idex HS), and focused on a PMT (PMT2101, Thorlabs). A 561nm picosecond-pulsed laser (BDS-SMY-561, Becker & Hickl) was power-modulated using a rotary neutral density filter (NDC-50C-4M-A, Thorlabs), reflected off a 573LP dichroic beamsplitter (FF573-Di01-25x36, IDEX), passed through the 552LP and 488LP dichroic beamsplitters, and focused into the patch cord. Red fluorescence exiting the patch cord passed through the 488, 552, and 561LP dichroic beamsplitters, was filtered using an interference filter (HHQ630/60, Nikon), and focused onto a PMT (PMT2101, Thorlabs).

The PMT signal and laser reference signal outputs were connected to the FLIPR analog-computing unit, which was connected to an NI data-acquisition system (NI-9261, NI 9215, and NI 9215, National Instruments) controlled by custom MATLAB software. Red and green lifetime signals were processed using separate analog computing units, DAQ boards, and computers, and were synchronized using a patterned voltage signal connecting the two DAQ systems. The key computation of the FLIPR analog-computing unit consists of the frequency mixing of the PMT signal with a phase-shifted reference signal (0, 0.5, 1, or 1.5π), resulting in a low-frequency signal (KHz) that is captured in the DAQ system. The lifetime signal is calculated using custom MATLAB software, using formula 2:

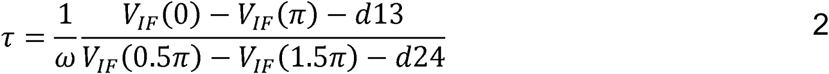

Wherein, τ is the fluorescence lifetime, V_IF_ is the intermediate frequency produced by the frequency mixers with phase-shifted reference signals (0, 0.5, 1, or 1.5π), and d13 and d24 are calibration factors compensating for voltage offsets in the system. FLIPR systems were calibrated using coumarin 6 (#442631, Aldrich) in ethanol for the green FLIPR system and rhodamine B (83689-1G, Sigma) in ethanol for the red FLIPR system. Further information on the components and computation of the FLIPR analog-computation unit is provided here.

### Optogenetic manipulation

Optogenetic manipulation was performed using a blue 473nm laser (MBL-III-473-100mW, Opto Engine LLC) and a red 593nm laser (MGL-F-593.5-100mW, Opto Engine LLC) for gtACR and ChrimsonR stimulation, respectively. Laser power was modulated using a programmable pulse generator (Master8, AMPI) in either TTL or analog mode. Laser power per setting was determined using an optical power meter (PM400, S120C, Thorlabs). Laser light power was calibrated to elicit physiologically relevant tonic and phasic dopamine changes of small, medium, and large magnitudes, as observed in the correlation experiments. Phasic activation (1.5s, continuous, red) and phasic inhibition (5s, continuous, blue) were repeated 10x in 3 different power settings within a session with an ISI of 2 min. Tonic activation (5 min, 20Hz, 30ms pulse width or 50Hz, 50ms pulse width, red) and tonic inhibition (5 min, continuous, blue) were repeated 3 times within a session, with 2-3 different power and/or frequency settings and an ISI of 5 min. Optogenetic stimulation was performed in a behavioral arena of 6.5 by 14.5 inches.

### Chemogenetic manipulation

Chemogenetic manipulation was calibrated through consecutive sessions of Deschloroclozapine (DCZ, HY-42110, MedChemExpress) injection at various concentrations with at least 48 hours in between sessions. During DCZ calibration, mice were first habituated for 5 minutes in an 8-by-8-inch behavioral arena. Mice were measured using FLIPR, and after 10 minutes, were injected with DCZ. DCZ dose calibration was performed to replicate physiologically relevant tonic dopamine change, as observed in previous behaviors, while minimally affecting phasic frequency and relative magnitude.

To compare the effect of DCZ on behaviorally evoked phasic dopamine release, mice were food-restricted overnight or to 90-95% of body weight and given approximately 10 chocolate pellets. Mice were measured using FLIPR, given 5 chocolate-flavored pellets (1 per minute), and subsequently injected with DCZ. After 10 minutes, the mice were given another 5 chocolate pellets (1 per minute).

For chemogenetic manipulation during progressive ratio experiments, mice were injected with saline or DCZ IP 15 minutes before the experiment. The order of receiving either saline or DCZ injections on subsequent days was counterbalanced such that half the animals received saline on day 1 and half of the animals received DCZ on day 1.

### Frequency analysis of photometry recordings

Frequency analysis of lifetime photometry recordings was performed using a fast Fourier transform (FFT) on the 20 Hz recording. For these frequency analyses, an entire two hour trace of fluorescence lifetime signal or a fully shuffled version of this signal was passed into the Fourier transform. Power at a given frequency was then quantified by taking the square of the absolute value of the transformed signal at any given point and then scaling this by the length of the entire FFT.

### Analysis of fluorescence lifetime signals during progressive ratio

As described above, animals were trained to operantly nosepoke towards an active port prior to being tested on the progressive ratio task. Fluorescence lifetime signals were initially acquired at 10 kHz and downsampled to 20 Hz for all analysis. Lifetime signals were aligned to behavioral data through video data and a synchronization pulse logged by Bonsai-Rx (v2.5.1). For analysis of phasic dopamine events relative to the reward-predicting cue, data was aligned to when the FED3.0 device reported that a pellet was dispensed. For analysis of changes in tonic dopamine during this task, tonic dopamine signals were extracted using a 1-minute moving median filter of the dopamine data. The change in tonic dopamine between reward and breakpoint was calculated as the change in the tonic dopamine level after reward was delivered (as an average signal of 2-4 minutes post break) until the break (as an average signal of 2 minutes before the break). Nosepokes were identified through logs generated by the FED3.0 device.

### CNN analysis of dopamine to PKA transformation

To map dopamine (FLIM-rdLight1.1, abbreviated as FLIM-rdLight here) dynamics onto PKA (FLIM-AKAR) dynamics, we trained a one-dimensional convolutional neural network (CNN) that predicts a FLIM-AKAR segment from a matched rdLight window of equal length. We used a 1 minute trace for both training and for predicting ahead given that this was roughly the timescale on which tonic information was assumed to be encoded. The rdLight and FLIM-AKAR data across all recordings was aggregated for training, validation, and testing.

Data was first divided per session into chunks each 20 minutes in duration. Then, half of these chunks were randomly set aside as held-out testing data (“held out data”) and not used for training or validation. Subsequently, all data chunks were aggregated and further subdivided into prediction FLIM-rdLight/target FLIM-AKAR pairs of 1 minute of FLIM-rdLight signal and a proceeding 10 seconds of FLIM-AKAR signal with the predictive FLIM-rdLight signal ending before the start of the FLIM-AKAR recording. This ensured the model was a purely predictive model rather than using current or future dopamine data to predict PKA activity. Pairs of recordings for training and validation were overlapping by at maximum 5 seconds given that the dominant frequency band for dopamine signaling was identified ∼0.2 Hz (Fig S1A). Finally, these input/target pairs were split into a training and validation set of 85%/15% five-fold times to perform k-fold cross-validation.

The training inputs and targets were then z-scored. The inputs were then passed into a network which was made up of a stack of six dilated temporal convolutions (kernel size 7; dilations 1, 2, 4, 8, 16, and 32; 64 hidden channels with ReLU nonlinearities), followed by a depthwise convolution that summarized the full input lookback window and a pointwise convolution that mapped the resulting features to the FLIM-AKAR prediction. The model was optimized with AdamW to minimize mean squared error (MSE) between predicted and observed FLIM-AKAR in the training set of 85% of data. Halting of the optimization and parameter tuning process was determined based on lack of changes in the MSE between the predicted and actual validation FLIM-AKAR data (constituting the other 15% of the dataset).

Following this train/test validation, the model was then tested on the held out data wherein the held out data was split again into input rdLight/target FLIM-AKAR pairs of the same length as in training/validation. This yielded several FLIM-AKAR traces with overlap given that subsequent rdLight traces were allowed to overlap in testing. The overlapping portions of the FLIM-AKAR signal were averaged, and the entire segment was output as a predicted FLIM-AKAR signal. The correlation between the actual and predicted FLIM-AKAR signal was then plotted in a representative trace as well as in scatterplot in Figure 3.

### Deconvolution analysis of lifetime signals

To determine the presence or absence of phasic signals, we used OASIS to determine the point-by-point probability of the occurrence of a phasic event. For determining the relationship between tonic dopamine levels and peak of dopamine phasic increases/decreases, the top 500 most likely phasic events were identified as putative events and used for analysis.

For determining the relationship between tonic dopamine and the probability of a PKA phasic event, deconvolution and phasic analysis was performed using OASIS. The trace was then normalized to the maximum probability returned by the OASIS algorithm across the entire session before regressing tonic dopamine level.

Finally, for determining the explanatory power of absolute versus phasic dopamine events for explaining PKA activity, first PKA events were identified using the OASIS deconvolution algorithm. For each PKA event, the preceding dopaminergic fluctuation was then extracted. This snapshot of a dopamine trace was then cross-correlated against the entire dopamine trace to identify 30 events which showed similar kinetics, albeit at different absolute levels, each with their own corresponding FLIM-AKAR trace. The sum of each of these 30 dopamine events was regressed against the sum of their corresponding FLIM-AKAR events, either taking the sum of the phasic trace (i.e. with the tonic component subtracted out) or the sum of the absolute trace. The correlation coefficients were compared using a Fisher r-to-z transformation and the resulting z-values were represented in the boxplot in Figures S5M,Q.

### Histology and Immunohistochemistry

Mice were anesthetized with high-concentration isoflurane. Mice were subsequently perfused transcardially with phosphate-buffered saline (PBS) and 4% paraformaldehyde (PFA) in PBS. Brains were extracted and stored in PFA overnight. Subsequently, brains were transferred in PBS until sliced. Brains were sliced on a vibratome (VT1000s, Leica) into 50-70µm slices. Slices were subsequently either mounted directly or stored for histology. Histology was performed by washing slices using PBS, then blocking slices at room temperature for 2 hours in 0.1% TritonX-100 /5% NGS (normal goat serum, Abcam) PBS. Slices were incubated in primary antibody solution (1:1000) overnight. Next, slices were washed using PBS with 0.1% TritonX-100 and incubated using secondary antibody solution (1:500) in blocking solution. Slices were subsequently washed using PBS with 0.1% TritonX-100 and stored in PBS before mounting. Slices were mounted on slides using ProLong Diamond Antifade Mountant with DAPI (ThermoFisher Scientific). Slides were imaged using a slide scanning fluorescence microscope (VS200, Olympus).

### Quantification and statistical analysis

Data analysis was performed using custom scripts in Python (v3.8) and MATLAB (v2021a and v2024b). Experiments were not blinded, and a pre-calculation of the statistical sample size was not performed. Two-sample t-test, paired-sample t-test, and regression ANOVA were applied as described in the Fig. caption. Single star (*), double star (**) and triple star (***) represent p<0.05, p<0.01 and p<0.001, respectively.

