## Supplemental Figures for "Tonic dopamine regulates cell-type-specific PKA activity and motivated behavior"

**Supplemental figures**
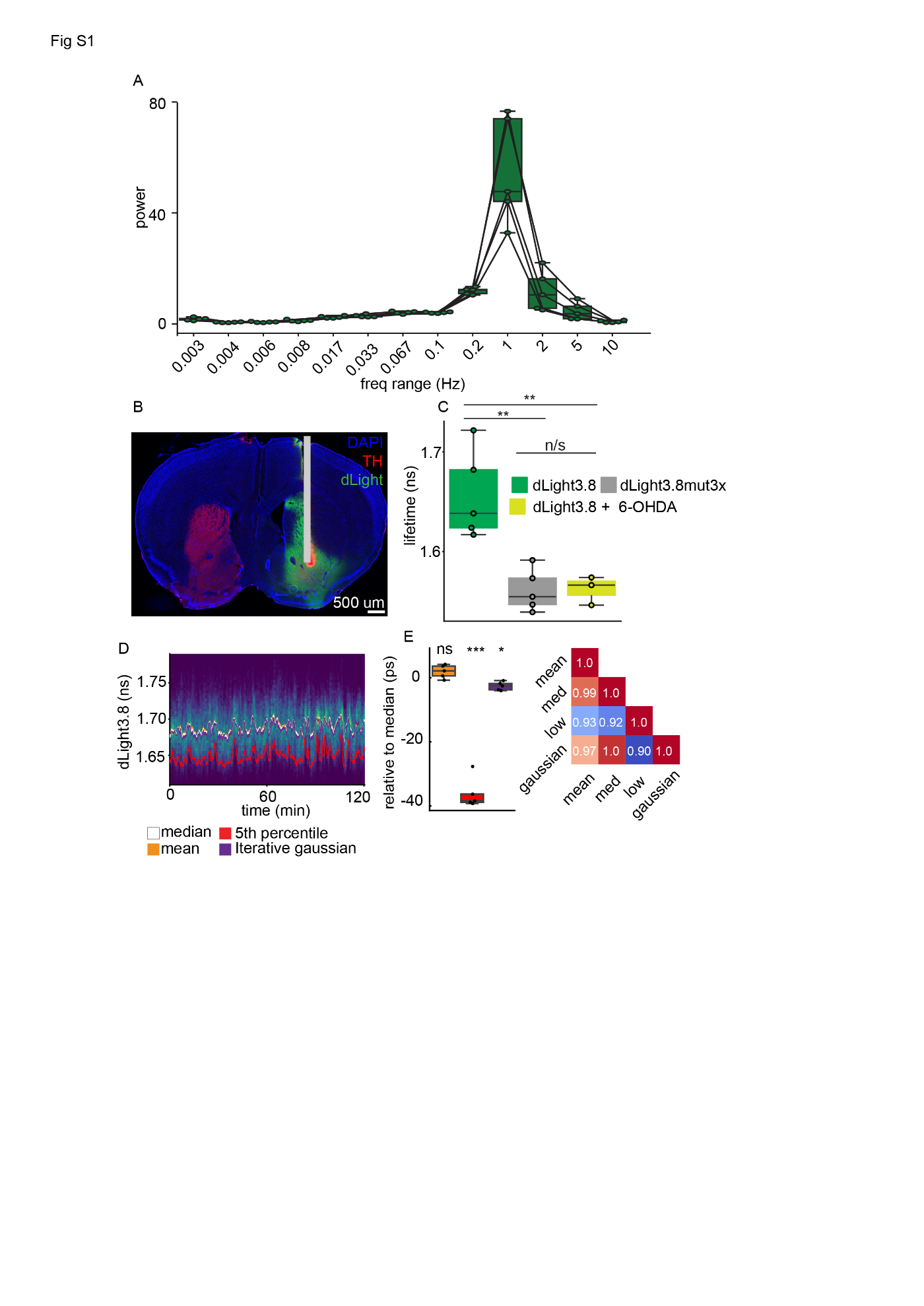

**Fig. S1:** Extended analysis of lifetime fluctuations in dLight3.8 and related sensors.

**A)** Comparison of power of lifetime signal in dLight3.8 in different frequency bands (n=5 mice). Each point represents power in the band between labeled frequency and next lower frequency.

**B)** Example histological section of animal injected with dLight3.8 (green) and ipsilaterally injected at the medial forebrain bundle with 6-OHDA for DAN (red) ablation and fiber optic implanted (gray bar).

**C)** Average lifetime measured in animals injected with dLight3.8 alone (n=5 mice), dLight3.8x3mut (n=5 mice) or dLight3.8 and 6-OHDA (n=3 mice) and comparison of these signals (dLight3.8 vs dLight3.8x3mut: p=0.002; dLight3.8 vs dLight3.8 + 6-OHDA: p=0.006; dLight3.8x3mut vs dLight3.8 + 6-OHDA: p=0.932; unpaired t-tests).

**D)** Example trace of lifetime measurement of dLight3.8 graphed as a pseudocolored rolling histogram with multiple extracted tonic dopamine traces overlaid. Data is the same as in Fig. 1.

**E)** Comparison of approximations of tonic dopamine to median dLight3.8 lifetime (left; mean: p=0.096, 5^th^ percentile: p=7.55x10^-5^, iterative gaussian: p=0.011; n=5 animals, one-sample t-test), and covariance matrix comparison of these approximations (right).

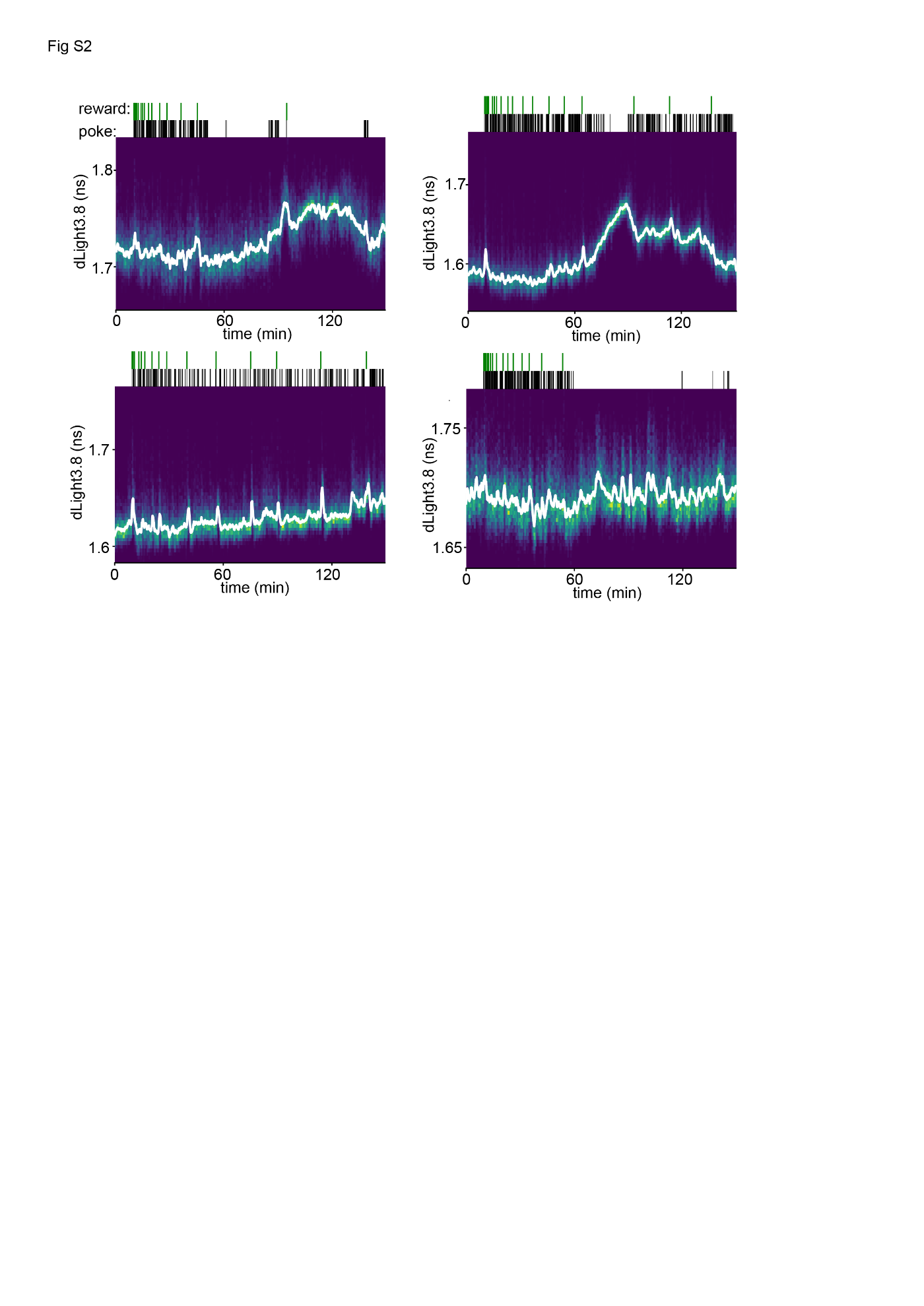

**Fig. S2:** Additional example traces of dLight3.8 lifetime fluctuations during progressive ratio task. Neural recording and behavioral data represented as in Fig. 2. Each plot represents a different animal and a different session during the progressive ratio task.

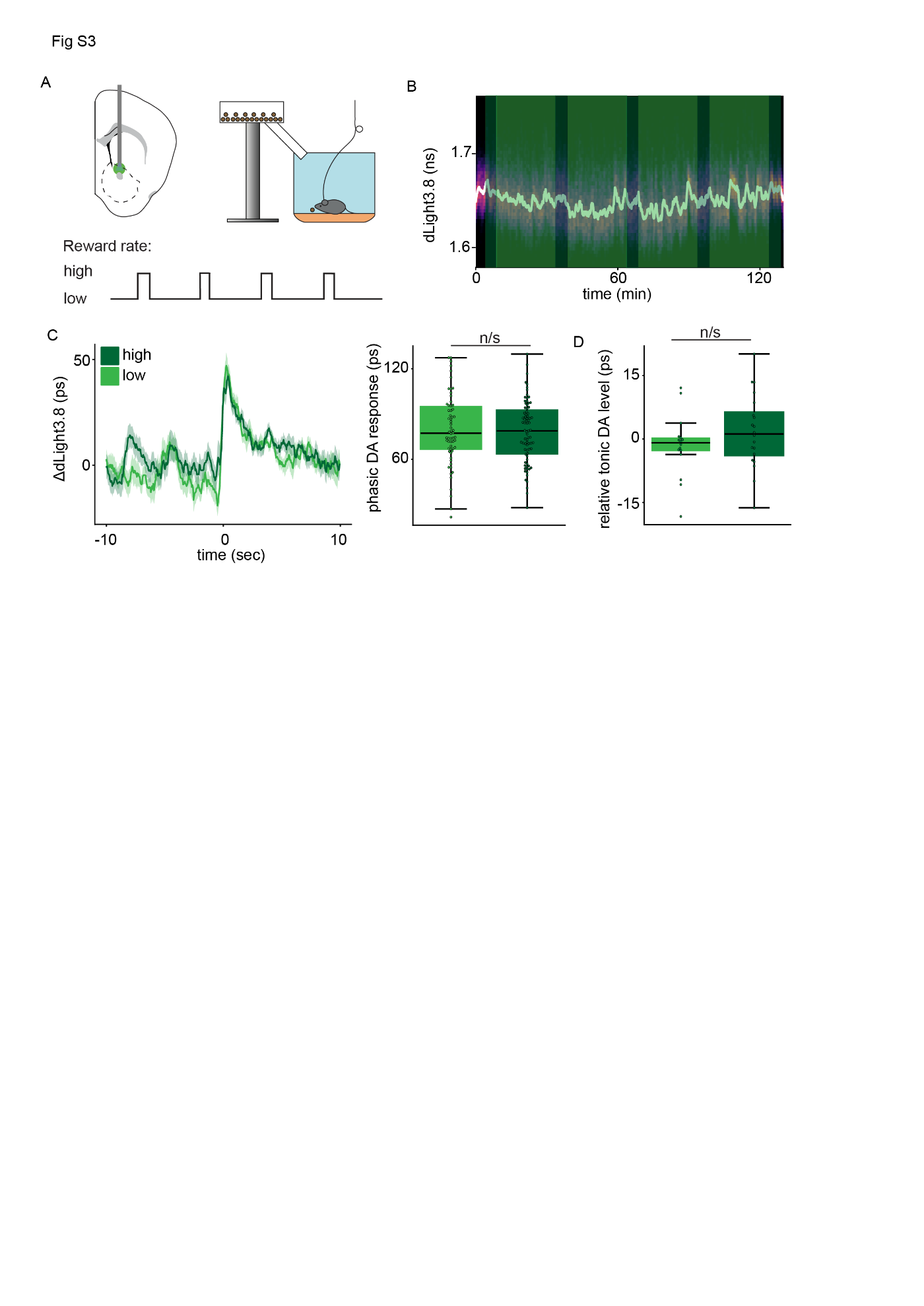

**Fig. S3:** Dopamine fluctuations during reward rate behavioral experiment

**A)** Schematic of surgical and behavioral protocol for measuring effects of differing reward rates on tonic dopamine levels.

**B)** Example of change in dLight3.8 lifetime during reward rate experiment. Dark green bar represents periods of high reward rate, light green bar represents periods of low reward rate.

**C)** Dopamine response to receipt of chocolate pellet during the high (dark green) and low (light green) reward rate period and quantification of these changes (n=62 pellets at low reward rate, n=79 pellets at high reward rate; p=0.743, unpaired t-test).

**D)** Session-normalized change in tonic dopamine during periods of high (n=20) and low (n=16) reward rates (p=0.236, unpaired t-test). For this set of experiments, n=4 mice used.

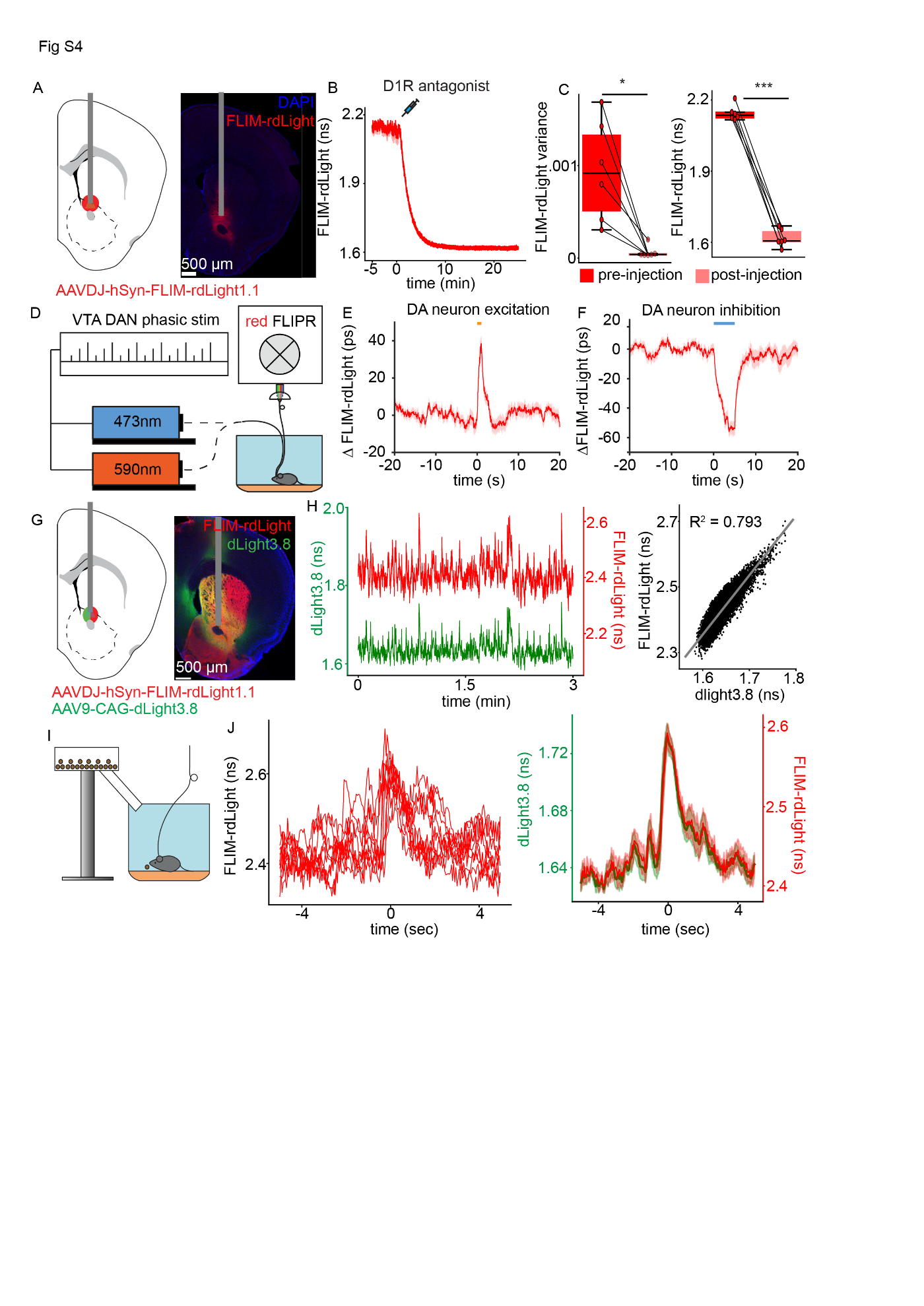

**Fig. S4:** Validation of dopamine-dependent changes in the fluorescence lifetime of FLIM-rdLight1.1

**A)** Schematic of injection/surgery in the NAC (left) and associated histological section of FLIM-rdLight1.1 sensor spread (red) and fiber placement (grey bar).

**B)** Average lifetime response of FLIM-rdLight1.1 sensor to intraperitoneal injection of D1R antagonist, SCH23390.

**C)** Change in variance (left; p=0.011, paired t-test) and absolute level (right; p=9.19x10^-7^, paired t-test) of rdLight lifetime signal (n=6 mice).

**D)** Schematic of optical setup used for simultaneous optogenetic manipulation of dopamine neurons and readout of changes in rdLight lifetime signal.

E) Average lifetime response of FLIM-rdLight1.1 to optogenetic stimulation of midbrain dopaminergic neurons (n=37 stimulations across n=3 sites in n=2 animals).

F) Average lifetime response of FLIM-rdLight1.1 to optogenetic inhibition of midbrain dopaminergic neurons (n=29 stimulations across n=3 sites in n=2 animals).

**G)** Schematic of injection/surgery in the NAC (left) and associated histological section (right) for simultaneous measurement of FLIM-rdLight1.1 (red) and dLight3.8 (green).

**H)** Example simultaneous recording of dLight3.8 (green) and FLIM-rdLight1.1 (red) and correlation between these two lifetime signals (right).

**I)** Behavioral schematic for evoking phasic dopamine responses in the NAC through random chocolate pellet rewards.

**J)** Example traces of fluctuations in FLIM-rdLight1.1 lifetime when animal receives chocolate pellet (left) and comparison of pellet-associated average fluctuation (right) in lifetime of FLIM-rdLight1.1 (red) and dLight3.8 (green). For two-color measurement experiments, n=6 mice.

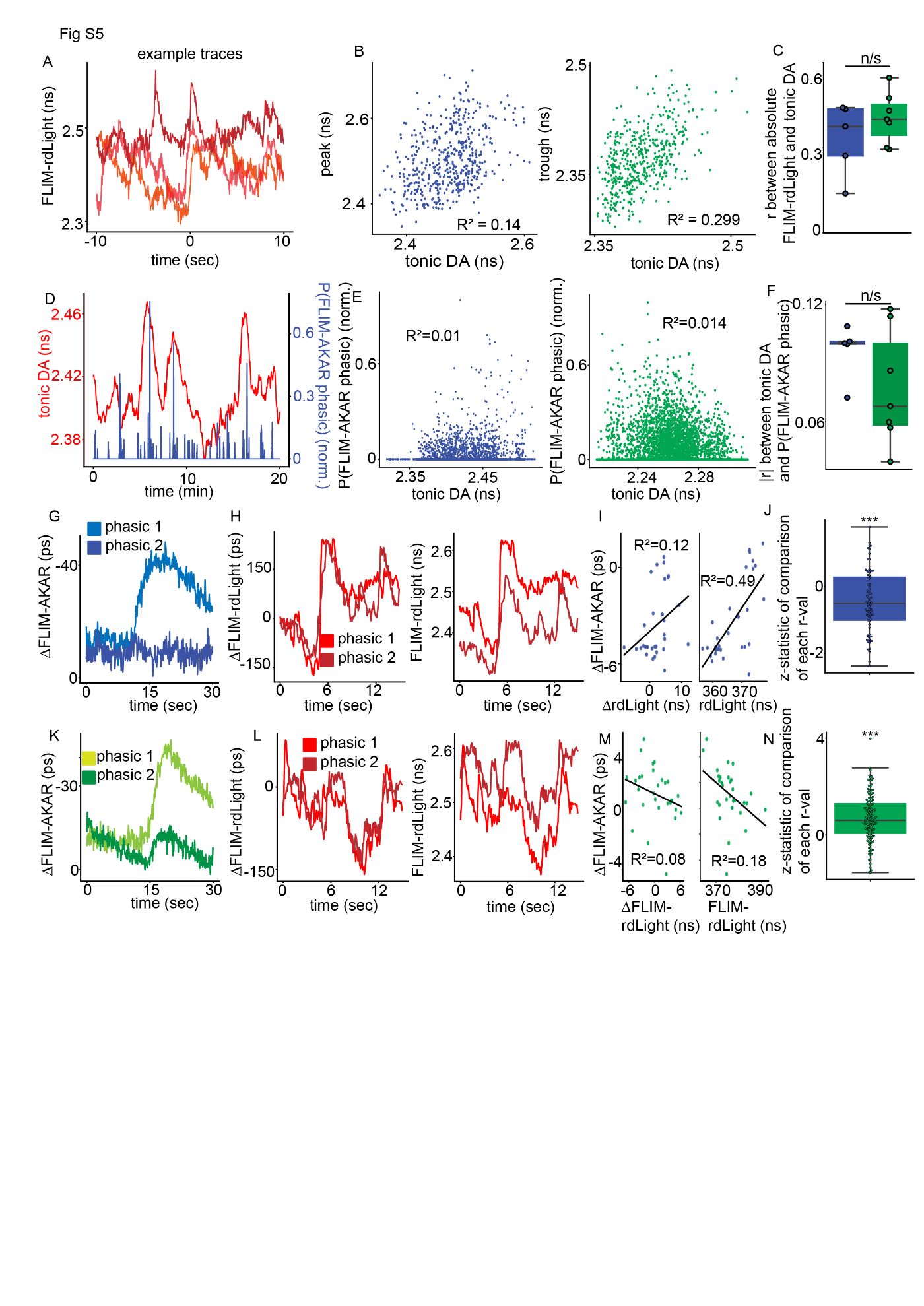

**Fig. S5:** Extended correlational analysis between dopamine and PKA signals

**A)** Example trace of spontaneously occurring phasic increases in dopamine at different tonic dopamine levels.

**B)** Correlation between tonic dopamine level and peak of a phasic dopamine increase (left) and trough of a phasic dopamine decrease (right).

**C)** Comparison of correlation between tonic dopamine level and peak/trough of phasic dopamine change in recordings of mice for D1R-SPN and D2R-SPN PKA dynamics (n=5 D1R-SPN animals, n=7 D2R-SPN animals; p=0.771, unpaired t-test).

**D)** Example trace of simultaneous changes in tonic dopamine level (red) and probability of a phasic D1R-SPN PKA event (blue).

**E)** Correlation between tonic dopamine level and probability of phasic PKA event in D1R-SPNs (left) and D2R-SPNs (right).

**F)** Comparison of correlation between tonic dopamine level and probability of phasic PKA events in D1R-SPNs and D2R-SPNs (n=5 D1R-SPN animals, n=7 D2R-SPN animals; p=0.173, unpaired t-test).

**G)** Two example traces of D1R-SPN FLIM-AKAR signal with different associated dopamine signals.

**H)** Dopamine signals associated with D1R-SPN FLIM-AKAR signals either normalized (left) or absolute (right).

**I)** Correlation between magnitude of D1R-SPN FLIM-AKAR signal and either normalized (left) or absolute (right) rdLight signal.

**J)** Boxplot of z-statistics for comparison of explanatory power (r-statistic) of the absolute vs normalized rdLight signal for D1R-SPN FLIM-AKAR dynamics (n=77 phasic changes, p=5.13x10^-5^, one-sample t-test).

**K-M)** As in panels G-I except for D2R-SPN FLIM-AKAR dynamics.

**N)** As in panel J except for D2R-SPN FLIM-AKAR dynamics (n=122 phasic changes, p=5.645x10^-14^).

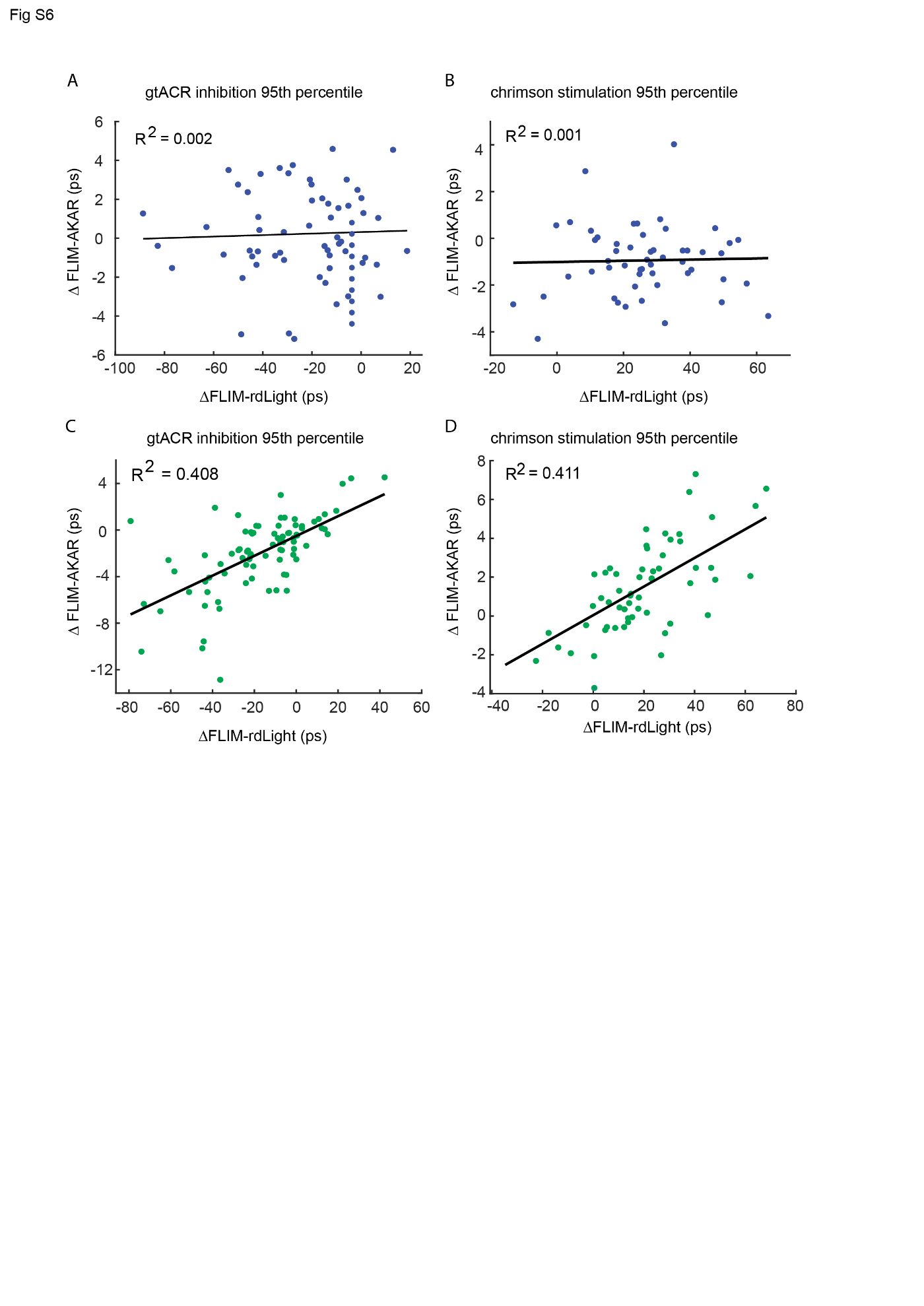

**FIG S6:** Correlation between change in optogenetically-evoked tonic dopamine and change in FLIM-AKAR signal when examining the 95^th^ percentile of the FLIM-AKAR signal.

**A)** Average optogenetically-induced tonic decrease in rdLight signal and associated D1R-SPN FLIM-AKAR dynamic, with regression between median of evoked FLIM-rdLight1.1 change (3-5min) and the associated 95^th^ percentile of FLIM-AKAR change in D1R-SPN

**B)** As in panel A except for changes seen with optogenetically-induced tonic increases in FLIM-rdLight1.1 signal.

**C)** As in panel A except in D2R-SPN FLIM-AKAR dynamics.

**D)** As in panel A except for changes seen with optogenetically-induced tonic increases in FLIM-rdLight1.1 signal and in D2R-SPN FLIM-AKAR dynamics.

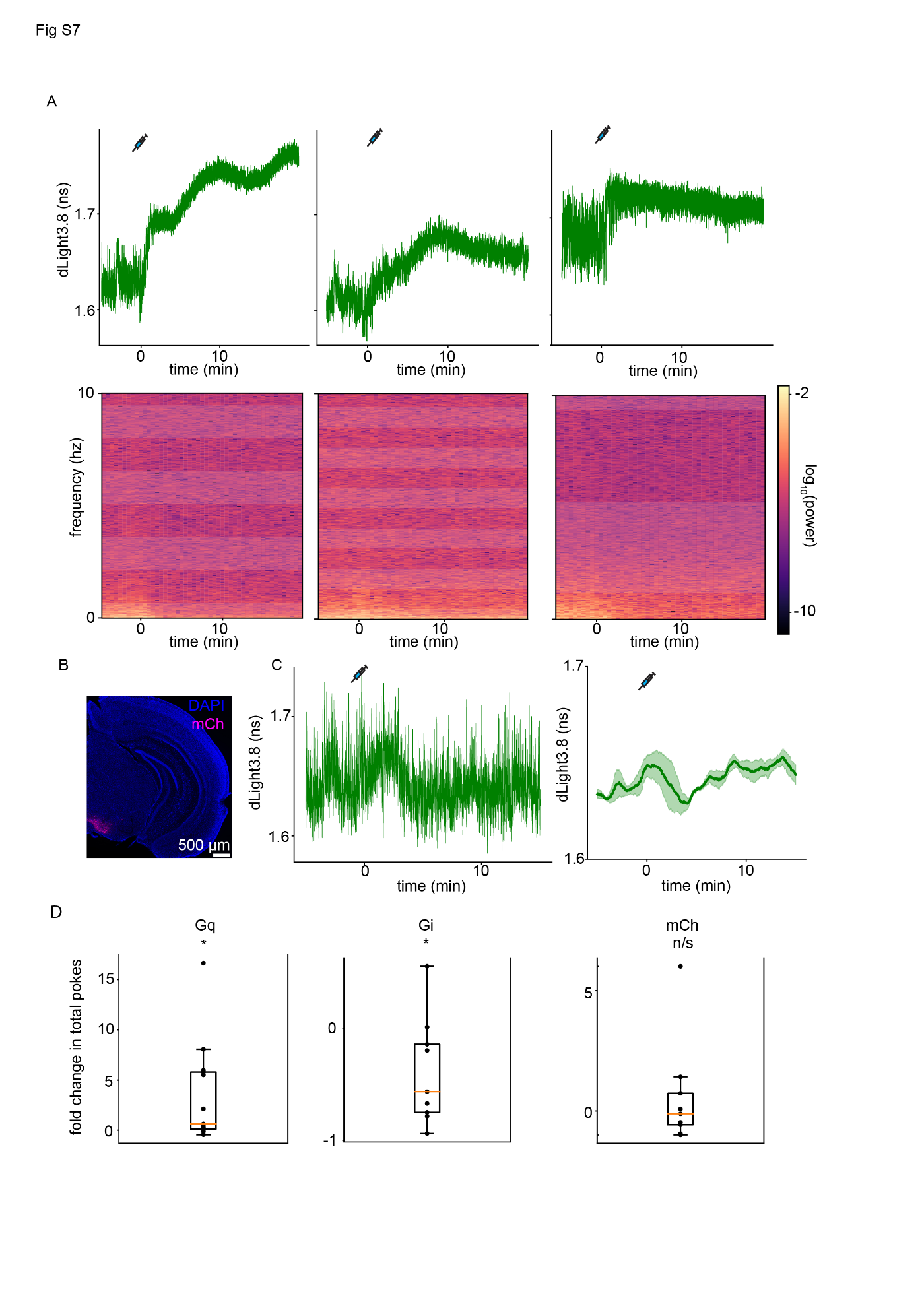

**Fig. S7:** Extended results of chemogenetic manipulation experiments

**A)** Example dLight3.8 lifetime responses to injection of 0.2 mg/kg of DCZ in animals expressing Gq DREADDs in VTA DANs (top) and associated spectrograms with each response (bottom).

**B)** Example histological section showing expression of mCherry in VTA DANs.

**C)** Example dLight3.8 lifetime fluctuations associated with injection of 1 mg/kg of DCZ in animals expressing mCherry in VTA DANs (left) and average lifetime fluctuation associated with this injection across mice (right). n=3 mice.

**D)** Fold change in number of pokes animals make prior to breaking when injected with DCZ vs saline (wherein a positive number corresponds to more pokes in the DCZ condition compared to the saline condition, and a negative number corresponds to more pokes in the saline condition compared to the DCZ condition) in Gq DREADD (n=13 animals, p=0.025; one-sample t-test), Gi DREADD (n=9 animals, p=0.042; one-sample t-test), and mCherry (n=9 animals, p=0.455; one-sample t-test) expressing animals.
